# Lattice Instability Drives Formation of Protofilament Clusters at the Microtubule Plus End Tips

**DOI:** 10.1101/2025.10.16.682920

**Authors:** Weizhi Xue, Jiangbo Wu, Tamara Bidone, Gregory A. Voth

## Abstract

Microtubules (MTs) are dynamic cytoskeletal filaments composed of α- and β-tubulin protein dimers. They are crucial for maintaining cell structure, facilitating intracellular transport, and ensuring proper chromosome segregation among other things. These biological functions are influenced by the dynamic instability of the MT plus-end tip. Recent simulations have discovered formation of protofilament (PF) clusters at the MT plus-end tip, but reliable extrapolation of PF cluster dynamics and detailed microscopic mechanism are yet to be developed to understand their behavior thoroughly. In this work, we have constructed, from “bottom up”, a relatively high resolution coarse-grained (CG) molecular dynamics (MD) model for tubulins with 20 CG sites per tubulin monomer, performed extensive CG MD simulations on MT lattices with 8 and 40 layers of heterodimers, and conducted comprehensive atomistic-level analysis. Our findings demonstrate that, in both GTP and GDP states, PF clusters are stable up to tens of microseconds of CG MD simulation time during spontaneous outward bending relaxation. PF clustering is initiated by longitudinal relaxation, stabilized by residual lateral interaction in the PF clusters. This process is thermodynamically driven by intrinsic lattice instability. In longer microtubules, this instability accumulates and further facilitates PF bending and clustering at the plus-end tip, but it can also be released via lattice curvature and supertwist. GDP-MTs form more PF clusters than GTP-MT on average and undergo more lateral cleavage and faster bending relaxation due to weaker lateral interactions, which facilitates MT catastrophe. GTP-MT forms flatter and more rigid PF clusters that favor nucleotide addition. Our findings highlight the critical role of lattice instability in microtubule dynamics and offer new insights on the conformational variability of MT plus-end tips.

**SIGNIFICANCE:** The dynamical structure and behavior of microtubule (MT) plus-end tips remains an important topic in structural and computational biophysics. For the recently discovered protofilament clusters at MT plus-end tips, this study provides a mechanism for their formation through comprehensive analyses of tubulin-tubulin interaction interfaces in all-atom simulations of complete MT lattices. This study also makes long-time predictions for the dynamics of PF clusters through a coarse-grained model for the tubulin heterodimers to extend the effective simulation time and system size, while preserving key details of the lateral and longitudinal interaction interfaces. The findings deepen understanding of the life cycle of PF clusters, while adding new insight on microtubule dynamics and the role of lattice destabilization.

## INTRODUCTION

Microtubules (MTs) are dynamic, cylindrical polymers composed of *α*- and *β*-tubulin heterodimers (**Fig. 1a**) that play a critical role in various cellular processes (1, 2). MTs are integral components of the cytoskeleton, providing structural support and maintaining cell shape (3, 4). MTs also facilitate intracellular transport, serving as tracks for the movement of organelles and vesicles as driven by motor proteins such as kinesin and dynein. MTs are furthermore crucial to cell motility and cell division, forming the mitotic spindle that ensures accurate segregation of chromosomes during mitosis and meiosis (2, 5). The *αβ* tubulin heterodimers are longitudinally aligned along the cylindrical axis to form protofilaments (PFs), while also interacting laterally to form a lattice with screw symmetry (6–9), as illustrated in **Fig. 1b**. The MTs usually contains one or more seam regions, where *β*-tubulin interacts with *α*-tubulin in the next PF and vice versa (9, 10). The highly versatile and dynamic functions of MTs arise at least in part from its dynamic instability, enabled by the nonequilibrium and dynamical structure at the MT plus-end tip. At that tip, MTs can then bind with other motor proteins and cellular organelles to perform biological functions (2, 11–14). MTs grow in length at the plus-end tip by binding of GTP-bound *αβ* heterodimers, known as the assembly phase. Through a lattice compaction (15), heterodimers undergo hydrolysis of GTP into GDP in the *β* subunit, and shrink through detachment of GDP-bound heterodimers from the MT-tip, known as the MT catastrophe (1, 2). Moreover, the plus-end tip has been experimentally observed to adopt multiple configurations, including sheet-like, curled, and flared structures, which all exhibit outward bending curvature, indicating the existence of lattice strain inside the MTs released at the plus-end tip (2).

**FIGURE 1.**
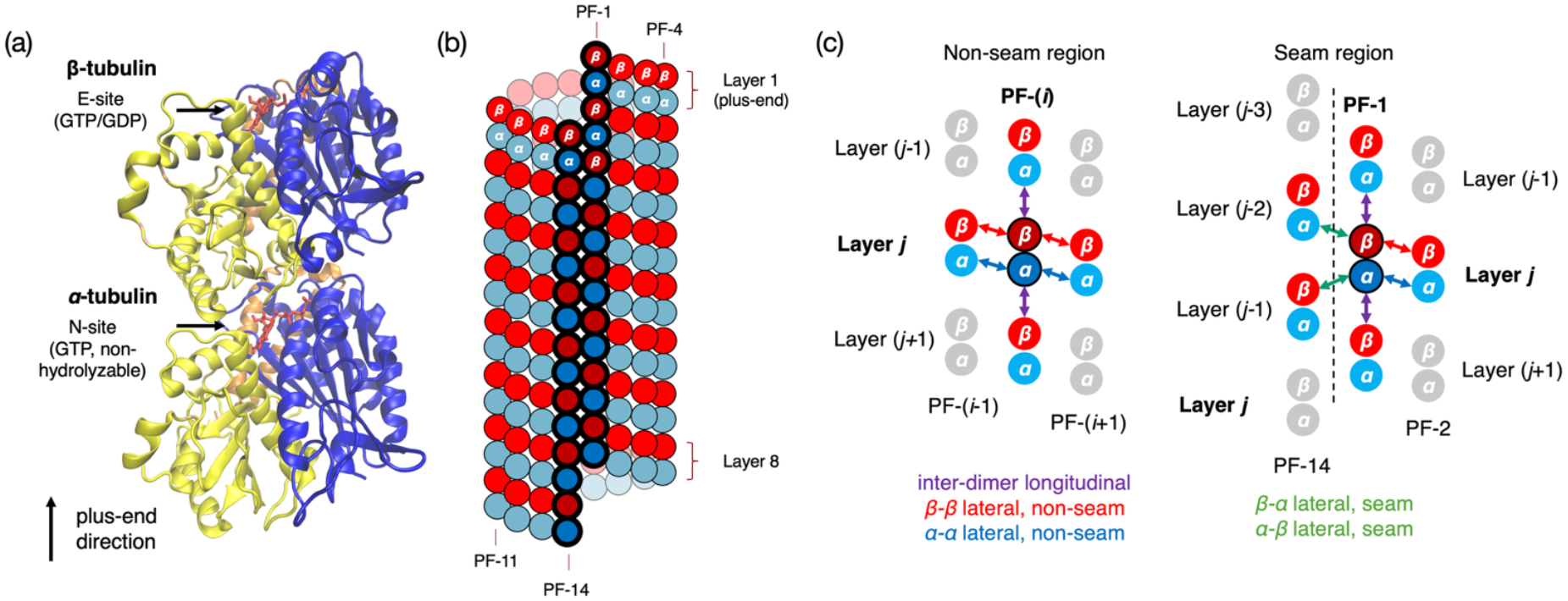
The structures of the tubulin heterodimer and the MT lattice. (a) The structure of *αβ* tubulin dimer. Each monomer contains a N-terminal domain (blue), an intermediate domain (yellow), and C-terminal domain (orange, in the back). The NTD is responsible for nucleotide binding, and only the nucleotide bound to *β*-tubulin is hydrolysable. (b) The structure of the MT lattice in the simulation of ref (16) and in the current study. It has 14 protofilaments and 8 heterodimer layers. The seam region is contoured in bold and darker colors. (c) Types of inter-dimer interactions in the MT lattice. In the methods section, only the nearest neighbors (colored) are extracted and aligned for forcefield construction.

The different behaviors of the MT tip during growth and catastrophe have long been attributed to the difference in lattice stability depending on the nucleotide state. So far, two models, namely the allosteric model (14, 17–19) and the lattice model (9, 20, 21), attempt to explain the dynamic instability of the MT tip based on the nucleotide state. The allosteric model hypothesizes that the dominant conformation of GDP-tubulin is more bent than that of GTP-tubulin, whereas the lattice model states that, while both heterodimer conformations are intrinsically bent in free heterodimers and single protofilaments, the stronger lateral interaction in GTP-bound MTs (GTP-MT) is able to “suture” the intrinsically unstable MT lattice together, and that hydrolysis of GTP into GDP weakens the lateral interaction, thereby triggering lateral dissociation and catastrophe. It should be noted that early all-atom molecular dynamics (MD) studies showed that GTP and GDP tubulin dimer were similarly bent (22), and that protofilaments constructed from those tubulin dimers exhibited essentially the same bending behavior (23). These simulation results thus favored the lattice model. Both models suggest that the plus-end tip of the GDP-MT is more flexible and can more readily laterally dissociate than the GTP-MT.

Recent experimental studies, including both *in vivo* and *in vitro* imaging (2, 24–26), have shown that both GTP-MT and GDP-MT exhibit outward bending behavior and similar curvature at the MT tip, hinting that the MT plus end tip structure is not as sensitive to the nucleotide state as once thought. Based on recent findings, several theoretical and computational models have been proposed, spanning from all-atom (AA) MD simulations to phenomenological kinetic models. Gudimchuk *et al*. developed a one-site Brownian dynamics model (25) with phenomenological parameters fitted to experimental data, and proposed that microtubule catastrophe can be induced by either weakened lateral bonds or more rigid longitudinal interactions. They also developed a kinetic model (27) to account for growth and shrinkage at curved MT tips enabled by infrequent straightening of PFs and attributed the source of dynamic instability to higher activation energy of lattice straightening in GDP-MTs. Although theoretically sound, these models rely on top-down experimental input for parameterization. Meanwhile, experimental structures are usually obtained via freezing and/or with inhibitors, so entropy effects at higher temperatures are missing (2).

Previous computational studies (22, 23, 28–32) have attempted to understand microtubule dynamical instability at the atomistic level through MD simulations of free tubulin heterodimers, single protofilaments, or lattice patches. Recently, with the ongoing growth in computational power, microsecond-level AA MD simulation has been possible for the MT lattice (16, 31), these have provided new structural insights from a “bottom-up” perspective (33–36). Igaev and Grubmüller have constructed an AA model (31) consisting of 14 PFs with 6 heterodimer layers and a helical rise of 1 tubulin (denoted as “14-1” MT, as in ref. (10)). They then simulated 5 replicas up to 2.4 µs, showing, clearly for the first time, that both GTP- and GDP-MT lattice tips exhibit spontaneous outward bending and lateral dissociation to form distinct protofilament clusters with residual lateral interactions. Based on their atomistic data and the latest experimental images, Kalutskii *et al*. were recently able to develop a molecular-mechanics CG model (32), at the resolution of 2 sites per tubulin, with direct cryo-ET observations of PF clusters to explain the formation of such PF clusters. Recent work by Wu *et al*. (16) has also reported such PF clusters in a 4 µs MD relaxation of an 8-layer 14-3 MT lattice. Not only are such PF clusters observed in the very early stage of the simulation, but they also appear stable regardless of nucleotide state. The GDP tip showed more clusters of fewer protofilaments each (on average), while the GTP tip had fewer clusters with more protofilaments in each, suggesting stronger lateral interactions in the latter case. With a machine-learning approach to learn the time evolution of coarse-grained radial coordinates, they were able to extrapolate PF cluster behavior up to 5.875 µs, roughly 2 times of the AA MD timescale. We note though that the actual timescale for MT tip relaxation spans much longer than a few microseconds, so the exact answer on the fate of these PF clusters still remains out of reach because of limits on computational power in AA MD sampling of such large systems (16).

How do these PF clusters form in the first place, and what is their eventual fate? In this work, we aim to bring our understanding of these questions to a deeper level than already accomplished in (16, 31). Specifically, we aim to address these specific questions: (a) How to reliably extrapolate the relaxation dynamics of these PF clusters? (b) What is the long-time behavior of these PF clusters in realistic MT tips, which have much longer lengths and possibly different numbers of protofilaments than simulations in refs (16, 31)? (c) What are the underlying microscopic structures, interactions and mechanisms that facilitate the formation of such PF clusters? For goals (a) and (b), we utilize the longest available 4 µs AA MD trajectories from ref. (16) as reference, and employ bottom-up coarse-graining (33–36) to the MT system, to build a CG model, in a higher resolution compared with refs (25, 32), to both extend simulation capacity and to capture key structural features in the tubulin longitudinal and lateral interactions. (In coarse-grained modeling, the term “resolution” refers to how many CG sites or “beads” are used to represent each amino acid or grouping of amino acids.) We then conduct extensive CG MD simulations to demonstrate that the MT plus end tips can form PF clusters stable up to tens of microseconds of CG MD simulation time (which is much longer than AA MD simulation time, as CG MD simulations are significantly accelerated) during the process of outward bending relaxation. The formation of the PF clusters is seen to be driven by an intrinsic thermodynamic instability of the MT lattice, with the GDP-MT lattice undergoing a more rapid PF cluster formation. We also construct a long MT model of 40 heterodimer layers and show that the accumulation of MT lattice strain further facilitates formation of PF clusters and outward bending, and the structural differences in nucleotide states are also observed to be more pronounced, in general agreement with experimental images in ref (32). For goal (c), we provide a detailed analysis of the longitudinal and lateral interfaces, identify key H-bond pairs responsible for the relaxation process from the all-atom MD data, and discover that the formation of PF clusters is initiated by relaxation of the longitudinal interaction and later reinforced and stabilized by lateral relaxation.

## METHODS

### Coarse-grained mapping of *α, β*-tubulins

To construct coarse-grained mappings for *α*- and *β*-tubulins, we applied the K-Means Clustering Coarse-graining (KMC-CG) method (37). This method is an extension of the earlier Essential Dynamics Coarse-graining (ED-CG) method (38) which overcomes the restraints enforced by sequence connectivity in the ED-CG approach and co-minimizing a new “loss function” combining the spatial vicinity of residues, while matching fluctuation modes and sequence integrity:

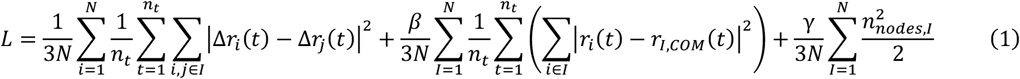

where *N* is the total number of CG sites, *n*_*t*_ is the number of frames in the MD trajectory of which the translational and rotational motion must be eliminated to a reference frame (“aligned”), the subscripts *i, j* denote the indices of C*α* atoms, the subscript *I* denote the indices of CG sites.

The reference trajectories in this work were obtained by extracting the C*α* atom trajectories of *β*- and *α*-tubulin from the AA MD simulation reported in previous work (16) separately and aligning them to their starting configuration. We used the *β*- and *α*-tubulins at the top of the microtubule lattice in PF-1 as they only have two neighbors (to their right and to the bottom, see **Fig. 1c**) and therefore are the least restrained. This approach allows more fluctuations in the loop regions and therefore allows for higher resolution in them than ED-CG or simple *N*-to-1 CG mappings. Then, a CG resolution of 20 beads per tubulin monomer was selected along with weight factors *β* = 1, *γ* = 100 to preserve more sequence integrity inside a CG site. We did not represent the nucleotide separately because the nucleotide state of the tubulin is already implicitly included in their equilibrium conformation and tubulin-tubulin interfaces. A total number of 50 optimization runs were calculated and the optimal CG mapping, rendered in **Fig. 2**, was determined as the mapping with the lowest *L*.

**FIGURE 2.**
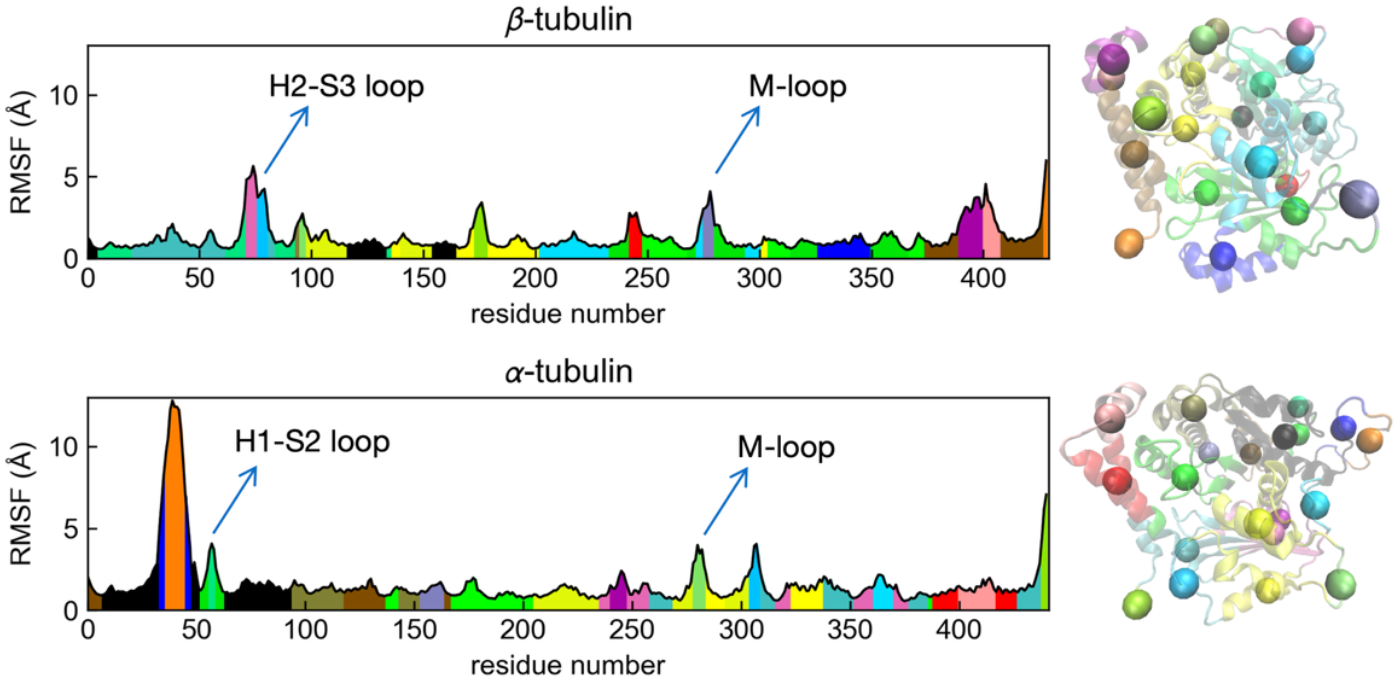
KMC-CG Mapping of *β*- and *α*-tubulins. Left: The RMSF from the reference trajectory is plotted against the residue index as the horizontal axis. The structurally relevant domains in lateral interaction, i.e., M-loop, H1-S2 loop and H2-S3 loop are marked with arrows. Right: The end point of the 4 µs configurations of the *β*- and *α*-tubulins are overlayed with the corresponding CG representations (spheres).

### CG forcefield parametrization

After obtaining the mapping schemes for *α*- and *β*-tubulins, we constructed the intra-dimer effective CG interaction bonds by applying the Heterogeneous Elastic Network Model (HeteroENM) (39). This model views the CG protein molecule as a network of effective harmonic springs within a cutoff distance, and is an effective small vibration approximation to the intramolecular forcefield without drastic conformational changes:

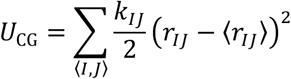

where *r*_*IJ*_ denotes the distance between a pair of CG sites within a given cutoff distance *r*_*c*_, and *k*_*IJ*_ is the corresponding spring constant.

From separate GTP-MT and GDP-MT trajectories reported in ref (16), we established intra-dimer and inter-dimer interactions for GTP-MT and GDP-MT separately and independently. The HeteroENM model requires that the reference trajectory be sampled at equilibrium to compute the ensemble average ⟨*r*_*IJ*_⟩, but according to previous work (16), such a system was shown to be unable to completely equilibrate in global degrees of freedom. On the other hand, local degrees of freedom, such as residue contacts on lateral and longitudinal interfaces (**Figs. 9-10 and Figs. S2.10-18**), typically relax on a faster timescale than collective motion. Hence, assuming a local-global timescale separation, we approximate the 3-4 µs segment of the AA MD trajectories as equilibrated in terms of local interactions, thus justifying the use of HeteroENM. For non-seam region, we extracted and aligned the trajectory of each heterodimer from layers 2 to 6 along with the 4 nearest neighbor heterodimers (see **Fig.1c**). For the seam region interactions only, we extracted and aligned each heterodimer in PF1 from layers 3 to 7, along with their 5 nearest neighbor heterodimers (see **Fig. 1c**). We then mapped these trajectories according to the KMC-CG mapping scheme and averaged the equilibrium bond lengths and spring constants for each interacting pair to ensure the microscopic symmetry of CG forcefield across different regions in the MT lattice. The cutoff of HeteroENM pairs was set to be 25 Å, in reference to the ~ 42 Å inter-monomer distance, to ensure that next nearest-neighbor interactions arising from coarse-graining are preserved while allowing sufficient intramolecular fluctuation. Pairs with *k* < 0.01 kcal/mol/Å^2^ were omitted. All HeteroENM calculations were performed using the OpenMSCG package (40). We directly used the averaged equilibrium bond lengths and spring constants from HeteroENM to construct intra-dimer harmonic bond interactions.

For inter-dimer longitudinal and lateral interactions, we used an *effective* Lennard-Jones 12-6 interaction as the CG bead interaction model. Taylor expansion of the LJ function at 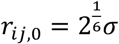 gives a small vibration approximation near the equilibrium pair distances, and the resulting coefficients, containing *ε*_*ij*_, and *σ*_*ij*_, should match *k*_*i j*,_ and *r*_*ij*,_ from HeteroENM directly, since HeteroENM is also a small vibration approximation. This gives the following approximation:

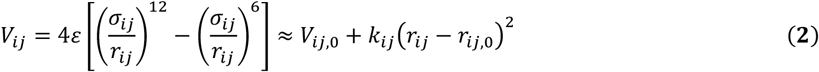

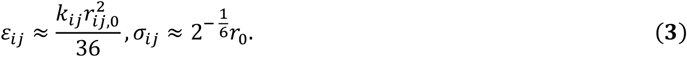

This approach circumvents the problem of insufficient sampling of protein-protein dissociation events, causing iterative CG forcefield optimization methods, such as relative entropy minimization (REM) and Iterative Boltzmann Inversion (IBI), to fail to converge to biologically realistic results, but instead allows for an approximate yet valuable and fully “bottom-up” parameterization for protein-protein interactions. The obtained forcefield parameters are available in the sample LAMMPS data file in Supporting Information, and comparison plots of the lateral and longitudinal interaction intensities are available in **Figs. S3, S4**.

### CG configuration generation

The AA MD trajectories was mapped with the OpenMSCG package (40) to directly obtain the starting configurations of the 8-layer model. To construct a 40-layer MT CG model, the starting configuration of the 8-layer model was translated in the direction of the MT axis (*z*-axis) and repeated. In accordance with observation from the AA trajectory (see **Fig. 10b, Fig. S2.20**), the distance between each heterodimer layer was increased by 0.5 Å in GTP-MT and 0.75 Å in GDP-MT, the least required to avoid lattice rupture due to release of lattice tension upon relaxation.

We also constructed a 13-3 CG model from the aforementioned 14-3 model, which conserved the longitudinal and lateral spacings as in the starting conformation. The tubulins were rotated with respect to the MT axis to correct the orientation. The distance between each heterodimer layer was increased by 0.25 Å in GTP-MT and 1.25 Å, the least required to avoid lattice rupture (see **Fig. S1.17** for benchmarks).

### CG MD simulation

All CG MD simulations of the 8-layer MTs were performed using LAMMPS version June 2022 and Apr 2025 (41), and were sampled in a constant *NVT* ensemble with a timestep of 10 fs under a Langevin thermostat (42) at 310 K after an initial energy minimization of 10^4^ steps. The starting configuration was chosen as the mapped starting frame from AA MD. The damping factor of the Langevin thermostat *τ* = 1/*γ* was set to be 1 ps by default and was varied among 0.1, 0.25, 0.5, and 0.75 ps to investigate its effects on the bending relaxation rate (discussed in Results and Discussions, see **Fig. 4e**). All CG sites of the bottom layer of heterodimers was placed under a positional restraint at their starting positions during minimization and simulation to mimic the effect of a longer MT lattice below the MT tip. When simulating the 40-layer MT model, positional restraint was applied to the bottom 3 layers at their starting positions, and the damping factor was 1 ps.

### All-atom MD data analysis

All atomistic trajectories from ref (16) were analyzed with the MDAnalysis package (43, 44) in Python except for secondary structure determination, for which we used the DSSP algorithm (45) implemented in the MDTraj package (46). Hydrogen bonds were determined with a cutoff of 4 Å and an angle cutoff of 30 degrees, for each longitudinal pair and lateral pair separately, and was averaged separately for different trajectory segments to determine H-bond frequencies (probabilities). We merged the frequencies of chemically equivalent H-bonds caused by equivalent atoms, e.g., terminal N atoms in Arg, terminal O atoms in Glu, Asp, and terminal H atoms in Arg, Lys. Lateral heterodimer pairs already dissociated in 3-4 µs were excluded for lateral interaction analysis, whereas all longitudinal pairs were considered in longitudinal interaction analysis due to no visible dissociation. To determine the leading H-bond interaction modes, we used Principal Component Analysis (PCA) and K-Means Clustering. We selected the top 20 most frequent H-bond pairs, averaged from all homolytic lateral pairs, from the 0-1 µs trajectory of the GTP-MT, and used a hyperbolic activation function 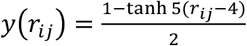 to build a 20-dimensional descriptor space. Data from all frames of all lateral pairs in both nucleotide states were combined to conduct PCA analysis, and we conducted K-Means clustering for cluster numbers 2 to 20 and eventually used 4 as the optimal cluster number and calculated the average descriptor vectors in each cluster.

## RESULTS AND DISCUSSION

### CG models of MT heterodimers

Figure 2. shows our KMC-CG mapping result for *β*- and *α*-tubulins with a resolution of 20 CG sites per monomer. According to ref. (9), the M-loop and H1-S2/H2-S3 loop regions are mainly responsible for the difference in lateral interaction between GTP- and GDP-MT lattices, characterized by larger distances of contact. Our CG mapping is therefore able to represent these loop regions with dedicated CG sites due to their higher root-mean-square fluctuation (RMSF), agreeing with physical insight. We have also tried lower CG resolutions 14 and 18, but under these resolutions, these key structural features may not be captured in both tubulins (**Figs. S1.1, S1.2**).

We compared the forcefields obtained in the Methods section for GTP-MT and GDP-MT in **Figs. S1.3** and **S1.4** for longitudinal and lateral interactions, respectively. From the *ε* parameters of their interactions, we observe that: (a) the longitudinal interactions in **Fig. S1.3** is stronger than lateral interactions in **Fig. S1.4**, as expected; (b) the effective lateral interaction in GTP-MTs are stronger than that in GDP-MT, especially for the M-loop – H1/S2 interactions and H9/H10 – H3 interactions (later analyzed in **Fig. 8**). According to the lattice model, such forcefields should lead to faster relaxation dynamics and more lateral dissociation behavior in GDP-MTs compared with GTP-MTs. This proposed trend is also in agreement with the reference all-atom MD trajectories in which the GTP-MT formed 4 PF clusters while GDP-MT formed 5 (Fig. 2). To further investigate this trend, we simulated the 8-layer CG model extensively (100 replicas) and compared the CG MD conformations with AA MD configurations (**Fig. 3**). The two most notable structural features in both AA and CG simulations are the outward bending motion and the formation of stable protofilament clusters.

**FIGURE 3.**
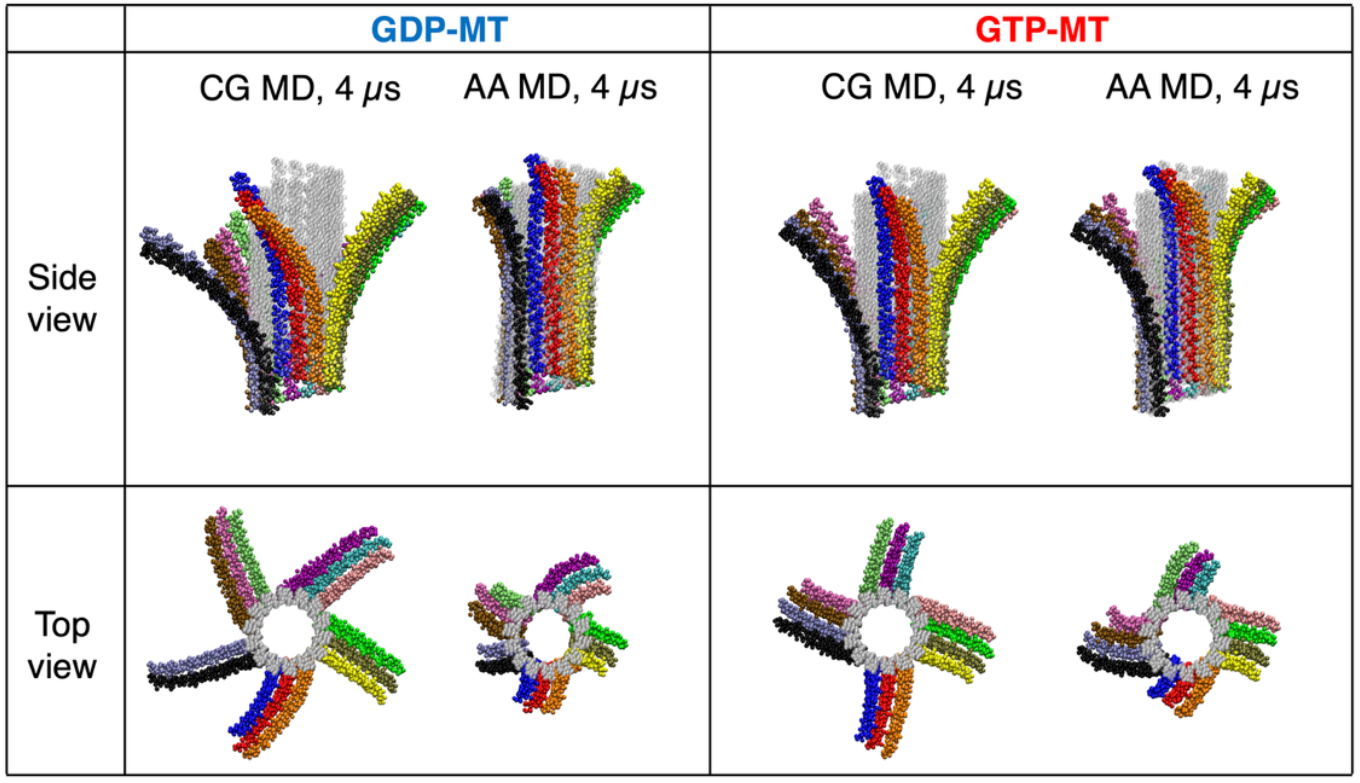
Comparison of the relaxed MT lattice at 4 µs, obtained from CG MD and mapped AA MD. The CG MD snapshots are selected among 100 independent 4 µs CG MD simulations, from which we observe a variety of PF cluster configurations (see **Fig. S1.7**). The AA MD snapshots are rendered from all-atom simulations in ref. (16). Colors represent different protofilaments, and the seam is between blue and black PFs (denoted as PF-1 and PF-14). The grey cylindrical configuration indicates the initial straight structure.

### Characteristics of bending relaxation in 8-layer MTs

We first investigated how quantify the extent of MT tip bending, we projected each tubulin in the *i*-th layer from top (*i* ∈ [1, 16]) of the *j*-th PF (*j* ∈ [1,14]) in cylindrical coordinates (*r*_*ij*_, *θ*_*ij*_, *z*_*ij*_,) illustrated in **Fig. 4a**, and plotted the radial profile *ρ*(*z*) of the PFs in **Fig. 4b**, relative to the center of mass of the bottom layer. We performed 100 replica CG simulations for both 14-PF and 13-PF models under the default CG forcefield and parameters and obtained the average radial profiles, and the results indicate that the plus-end tip of GDP-MT achieves a higher outward bending curvature than that of GTP-MT at 4 µs. We also started from the mapped AA configuration at 4 µs and performed 4 CG MD simulation replicas for another 20 µs (later referred to as “continuation runs”), and similar average radial profiles were obtained, indicating that GDP-MT exhibits more bending in the 8-layer model.

**FIGURE 4.**
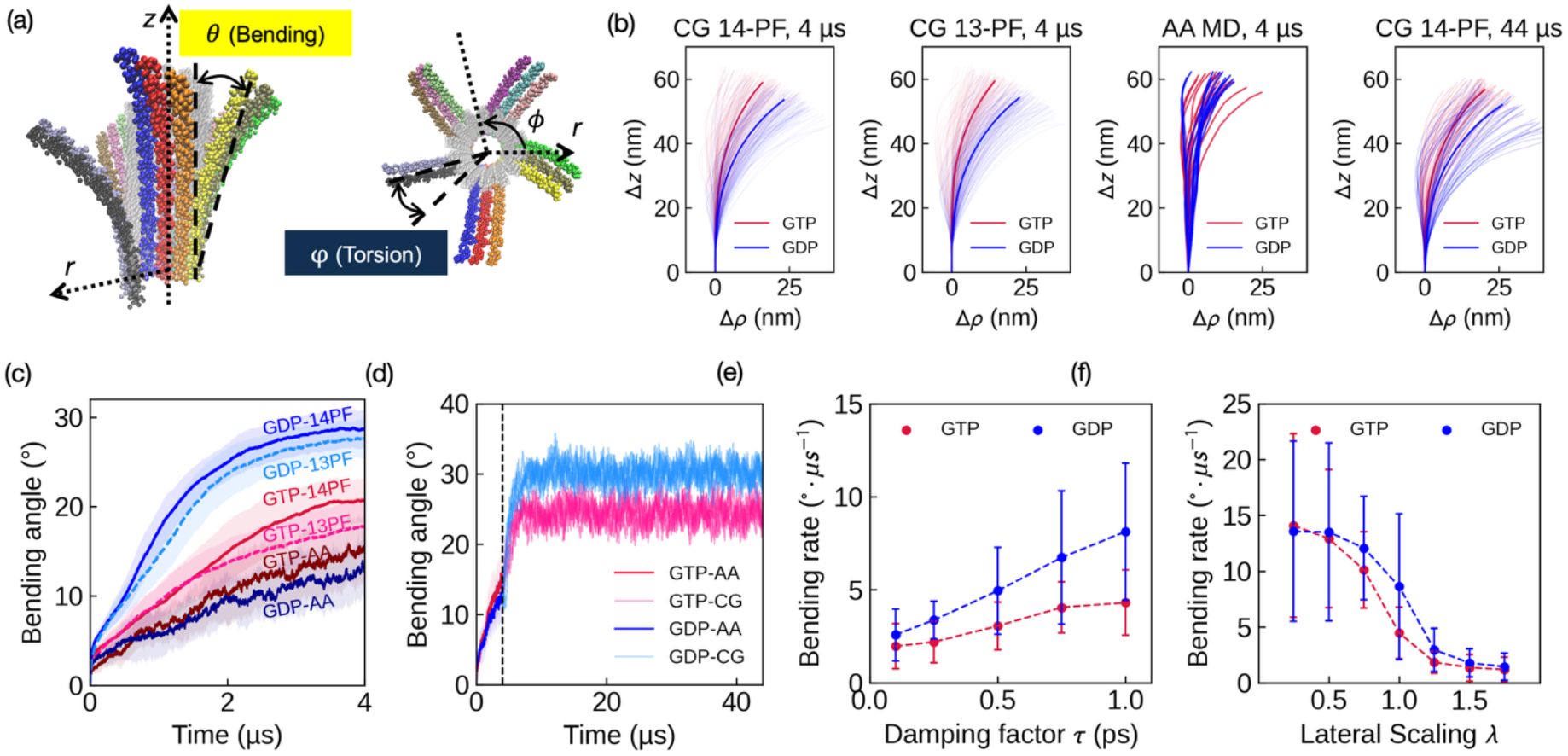
Bending relaxation of an 8-layer 14-PF microtubule lattice. (a) Definition of bending and torsion angles for each heterodimer. Examples are drawn for the top layer of the yellow protofilament (PF-4), and the top layer of the black heterodimer (PF-14). (b) Comparison of radial projection of the protofilaments in 14-PF CG frames at 4 µs (10 replicas), 13-PF CG frames at 4 µs (10 replicas), 4 µs AA MD frames, and CG MD configurations at 24 µs (5 replicas). (c) Comparison between bending curves: AA, 14-PF CG model and 13-PF CG model, for GTP and GDP. Both 14-PF and 13-PF CG bending curves are averaged from 100 replica simulations. Shaded areas indicate the standard deviation of the bending angle trajectories from the 14 individual PFs. (d) Comparison between CG MD bending curves, starting from 4 µs mapped AA configuration, plotted for 4 replicas. The first 4 µs represents the bending curve of the AA simulation. (e) PF bending rate as a function of Langevin damping factor τ. The standard deviation is computed from 10 replicas for each data point. (f) PF bending rate as a function of lateral interaction scaling factor λ. The standard deviation is computed from 20 replica simulations for each data point.

Alternatively, we define the bending angle for tubulin (*i, j*) as the angle between the MT axis vector and the vector pointing from the bottom layer heterodimer to tubulin (*i, j*), as illustrated in **Fig. 4a**. We calculated the bending angles for the 100 replicas and compared the time evolution of the bending angle between the AA model and CG model. There is a significant rate difference between the bending relaxation rate of GDP-MT and GTP-MT in CG simulations, although the AA MD simulation does not predict such a pattern due to limited number of replica available. The simulations of 13-PF models also agreed generally with 14-PF models, although the 13-PF GTP-MT bending angles are slightly lower near 4 µs than the 14-PF model. It is also reasonable that CG simulation achieves faster bending dynamics due to the fact that CG dynamics is accelerated (35). We also calculated the bending angles for the continuation runs, and from **Fig. 4d**, both GTP-MT and GDP-MT tend to stabilize and fluctuate at an “equilibrium” after less than 4 µs of CG MD simulation time, but the equilibrium bending angle of GDP-MT is higher than that of GTP-MT with statistical significance. Note that each 1 µs of CG MD simulation time approximately equals 2.5-3 µs of AA MD simulation time if we scale the MT bending rate of the former by the latter.

The lattice model suggests that the weakening of lateral contacts, together with the accumulated longitudinal strain, is the source of instability and outward splaying for GDP-MT lattice. To investigate this proposal, we varied the strength of lateral interaction by multiplying a scaling factor *λ* to the lateral interaction CG forcefields and performed 20 replica CG MD simulations for each value of *λ*, and so obtained the corresponding averaged bending curves (**Fig. S1.6**). From these bending curves we employed linear fitting to the linear growth ranges of the bending curves to extract the bending rates under different strengths of the lateral interactions. The results in **Fig. 4f** show a sigmoid-like decrease in bending relaxation rate as lateral interaction is strengthened. Moreover, the bending rate of GDP-MT lattice is larger than that of GTP-MT. This agrees with insight from our CG model parameterization that some strongest lateral pair interactions in GDP-MT have smaller *ε* parameters than those in GTP-MT. We also altered the Langevin damping factor of the CG simulation in the range of 0.1 to 1 ps and found that the relaxation rate increases linearly with the damping factor (**Figs. 4e, S1.5**), while the ratio between GTP/GDP relaxation rates stays roughly the same. This behavior indicates that the temporal evolution of the bending angle can be well modeled by Langevin dynamics in the overdamped regime. This result also suggests that altering the ionic strength and solute concentration could possibly regulate MT dynamics, as slower tip relaxation rates are associated with higher dynamical stability of the MT lattice. In short, our results reveal that the rate of bending relaxation is mainly determined by the strength of lateral interaction and dynamical friction, and the bending dynamics of the GDP-MT is indeed faster than that of the GTP-MT given the same simulation setup.

We have also noted that some PFs, particularly those in the left of a PF cluster, have a slightly inward bending tendency. As can be seen from **Fig. 4b**, there is coexistence between PFs bending inward and PFs bending outward, in the lower layers in both GTP- and GDP-MTs, indicating slight deformation to the cylindrical shape, possibly into an eclipse. Such coexistence suggests that the lower layers, although closer to the restrained layer and laterally bound, also have a tendency to experience lattice deformation. It is unclear how further this deformation propagates in a real, unrestrained MT lattice at room temperature, because (a) all experimental images of MT tips to our knowledge are observed at low temperature under which the bending dynamics of MTs are frozen and less entropy-driven, higher lattice symmetry is favored, and (b) it is computationally very expensive to simulate a longer MT lattice model with AA MD. These considerations make it necessary to carry out CG simulation on a longer MT lattice, as we have done in the next section.

### Number and configurations of PF clusters in the 8-layer model

In the all-atom MD simulations, both GTP- and GDP-MTs are observed to stabilize into PF clusters, with different configurations. We denote the different configurations with their breaking points, i.e., the PF index *i* where interacting neighboring PF-*i* and PF-(*i* + 1) dissociate (Fig. 5a), and the all-atom configuration is {3, 7, 10, 14} for GTP-MT and {3, 6, 9, 12, 14} for GDP-MT. Igaev and Grubmüller performed 5 independent simulations on 6-layer MTs (31) and found that these PF clusters are stochastic, while GDP-MTs tend to form slightly more splayed clusters than GTP-MT. We analyzed the cluster configurations in all 100 CG MD replicas under default forcefields and parameters and indeed found a various range of possible PF cluster configurations (**Fig. S1.7**), from which we could recover AA cluster configurations. From these simulations, we could calculate the average number of clusters formed in GTP-MT and GDP-MT, which are (3.93 ± 0.57) and (5.36 ± 0.62), respectively, and GDP-MT had a higher tendency to form 5 clusters than GTP-MT, as shown in Fig. 5b. In 13-PF simulations, these numbers are (3.39 ± 0.65) and (5.00 ± 0.62), respectively, showing the same trend as 14-PF simulations but with fewer PF clusters formed on average. We therefore hypothesize the GTP cap in the 13-PF model may have better stability overall when compared with 14-PF model, due to fewer PF clusters formed and slightly slower outward bending tendency.

**FIGURE 5.**
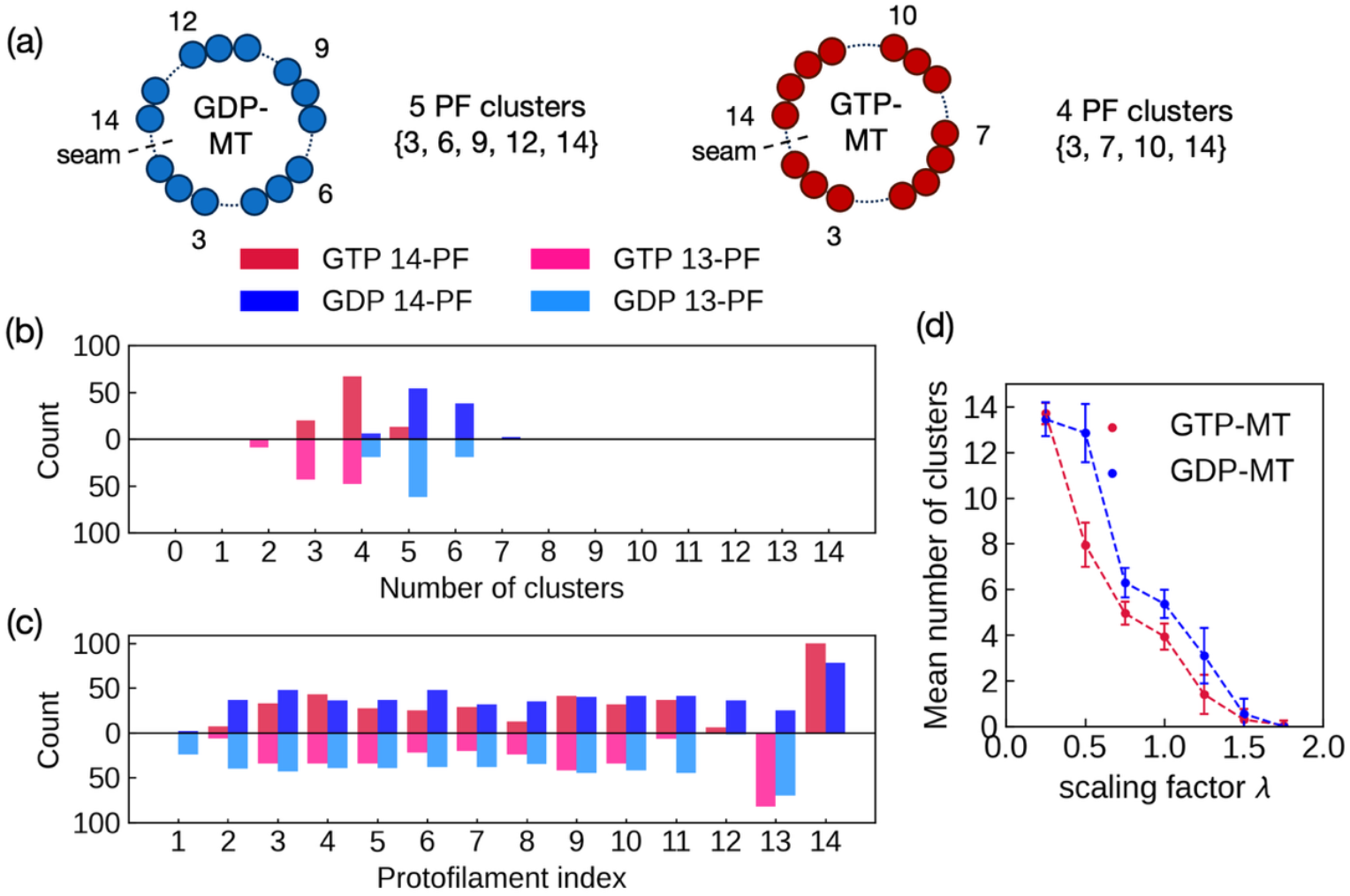
Distribution of tip cluster numbers in an 8-layer 14-PF microtubule. (a) The nomenclature used in this work to describe the configurations of the PF clusters. (b) Distribution of tip cluster number of the 100 replicas in both 14-PF and 13-PF models, each run for 4 µs. (c) Distribution of breaking points in both 14-PF and 13-PF models, i.e., the PF index ***i*** where interacting neighboring PF-***i*** and PF-(***i*** + **1**) dissociate, of the 100 replicas. (d) Mean tip cluster number computed from simulation (scatters) and approximative prediction (lines) as a function of lateral interaction strength in the 14-PF model. The standard deviations for simulation results are computed for 20 replicas except for ***λ*** = **1**, where they are computed for all 100 replicas. Note: it is our convention in this paper that the number of PF clusters includes single, separate protofilaments to differentiate the two limiting states, i.e., fully splayed (PF cluster number = 14) and lattice (PF cluster number = 0) states.

Additionally, we analyzed the average number of PF clusters and their standard deviations for simulations under different lateral interaction strengths, and the results are shown in **Fig. 5d**. It is the convention in this paper that the number of PF clusters includes single, separate protofilaments to differentiate the two limiting states, i.e., fully splayed (PF cluster number = 14) and lattice (PF cluster number = 0) states. Similar to the trend in the bending rates, there was a monotonous decrease in the average number of PF clusters, indicating that weakening of lateral interaction leads to formation of more PF clusters, i.e., the PFs at the tip are more likely to undergo lateral cleavage. This further supports the key hypothesis from the lattice model that catastrophe is triggered by weaker lateral interaction in GDP-MTs.

**Figure 5c** depicts the distributions of breaking points for both GTP- and GDP-MT for 14-PF simulations. The most thermodynamically probable dissociation is the seam region, i.e., between PF-14 and PF-1 indicates that the PF region is a constant opening for the microtubule lattice. However, we emphasize that this thermodynamic trend does not guarantee faster bending dynamics at the seam region, and the seam region can remain closed in a small portion of cases (see **Fig. 5c** and Fig. **S1.18**). Additionally, for GTP-MT, PFs 3-5 and 9-11 are slightly more favorable than their neighboring PFs. This trend is also seen in 13-PF simulations (**Fig. S1.18**), in which PFs 3-5 and 9-10 are more favorable.

In all 4 µs replica simulations of 14-PF and 13-PF models, the configurations of PF clusters tend to stabilize. In the 5 continuation runs, the PF clusters also remained stable with no further cleavage events, and the PFs in each cluster undergo collective bending and torsional movements, and their configurations also remained stable (See Supporting Movies).

### Bending relaxation and PF cluster behavior in 40-layer MTs

What is the bending behavior like in realistic MTs, with more layers and longer relaxation timescales, and different number of PFs? What is the fate of the PF clusters in these longer MTs? To investigate this, we constructed a 40-layer microtubule model and simulated with the baseline CG forcefield and a Langevin damping factor of 1 ps to accelerate the relaxation dynamics. All **10 independent** replica simulations were continued for at least 20 µs of CG MD simulation time (and again we note that this corresponds to a much longer real time). We have also conducted 10 independent simulations for 40-layer 13-PF models. **Figure 6** shows the configurations obtained from the last frame of these simulations, and we immediately notice several important structural features:

**FIGURE 6.**
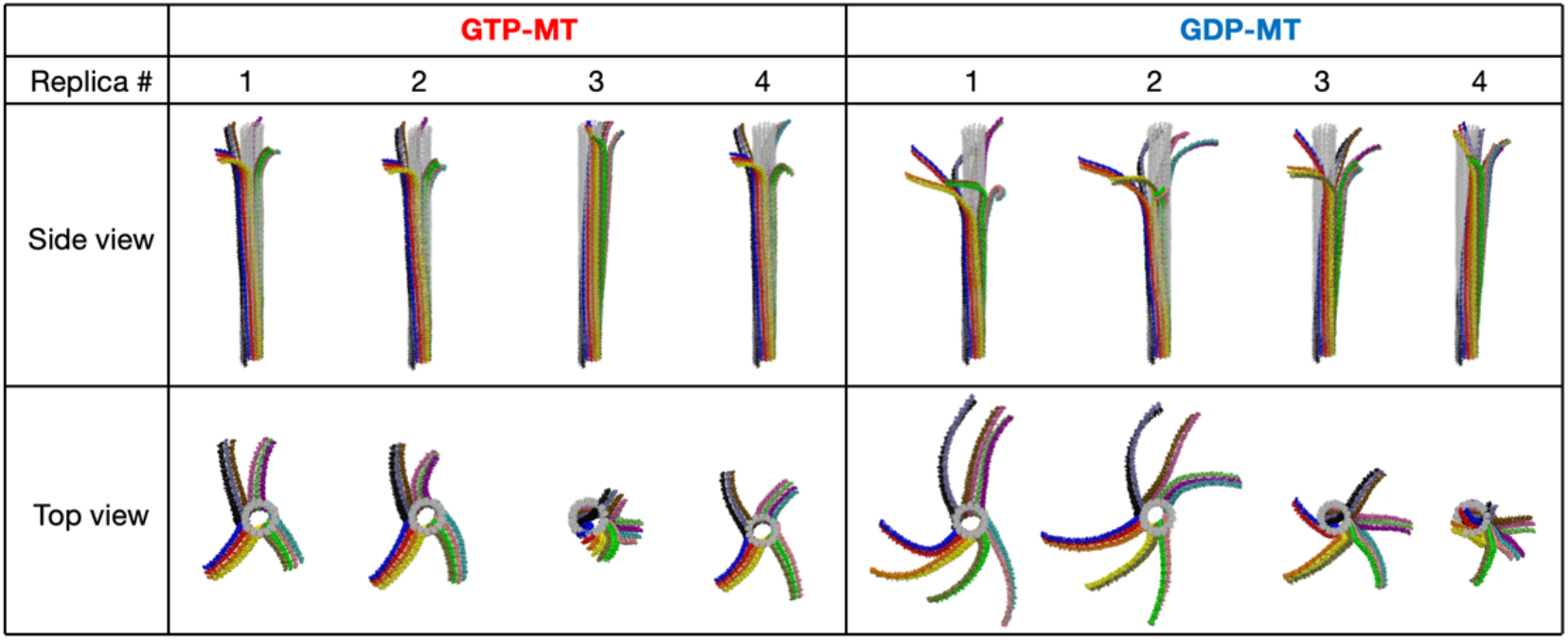
CG MD simulation snapshots on 40-layer 14-PF MT models. Snapshots at the end of simulation of 4 (out of 13 total) replicas of the 40-layer MT trajectory, each spanning at least 20 µs. Same color coding for different PFs as in Figures 2, 3. For all 13 independent simulations of the 14-PF model, see **Fig. S1.12**. For 13-PF MT snapshots, see **Fig. S1.13**.

#### (1) More PF clusters in GDP-MT

The PFs remain clustered in agreement with the average number of clusters obtained in 8-layer models. According to **Fig. S1.12**, of the 13 simulations of the 14-PF model, GTP-MTs and GDP-MTs form (3.92 ± 0.27) and (5.31 ± 0.46) PF clusters on average, showing a significant difference in the number of PF clusters. In 13-PF simulations, of 10 simulations, GTP-MTs and GDP-MTs form (3.6 ± 0.9) and (5.1 ± 0.3) PF clusters on average (**Fig. S1.13**). Moreover, the PF cluster configurations in GTP-MTs remained stable, but in GDP-MTs, some larger PF clusters were observed to undergo subsequent lateral dissociation into smaller clusters after their initial formation and stabilization.

#### (2) Continued, stabilized, and reversible peeling

We defined a lateral interaction pair to be dissociated if their center-of-mass (COMs) distance exceed 60 Å (which is approximately 1.5 times of their average COM distance in the lattice state). For each timestep in the trajectory, we calculated the number of heterodimer layers that underwent lateral dissociation, or “peeling”, and the results are shown in **Fig. 7a**. In both GTP-MT and GDP-MT, the number of layers that underwent lateral dissociation to form PF clusters (*n*_*p*_) continued to grow in 3 replicas, and the growth of *n*_*p*_ was more significant than GTP-MT. For the 19-20 µs segment of simulations in the 14-PF system, GDP-MT has (10.13 ± 2.87) layers peeled on average, whereas GTP-MT has (7.68 ± 2.26) layers peeled on average. This aligns with our expectation from the 8-layer simulation that: (a) the GDP-MT lattice is less stable and undergoes more lateral dissociation; (b) the system will try to apply the “learned” CG equilibrium distribution to anywhere in the lattice via constant peeling. This trend also applies to 13-PF simulations, but with less difference:GDP-MT has (10.17 ± 1.26) layers peeled, and GTP-MT has (8.57 ± 2.60), among 10 replicas. We also notice that, although the outward peeling motion appears thermodynamically favored, it is reversible, as some replicas in both GTP-MT and GDP-MT exhibited brief decreases in **Fig. 7a**, and one 13-PF simulation of GTP-MT exhibited near-total suturing of peeled layers **(Fig. S1.14)**. This reversibility is also present in as seen from **Fig. S1.15** if we look at the lateral openings between individual protofilaments. We notice that such peeling may become stabilized by lattice tilting, PF supertwist and elliptical deformation of tubulin layers, as seen from some plateauing behavior in **Fig. 7a** and **Fig S1.15**.

**FIGURE 7.**
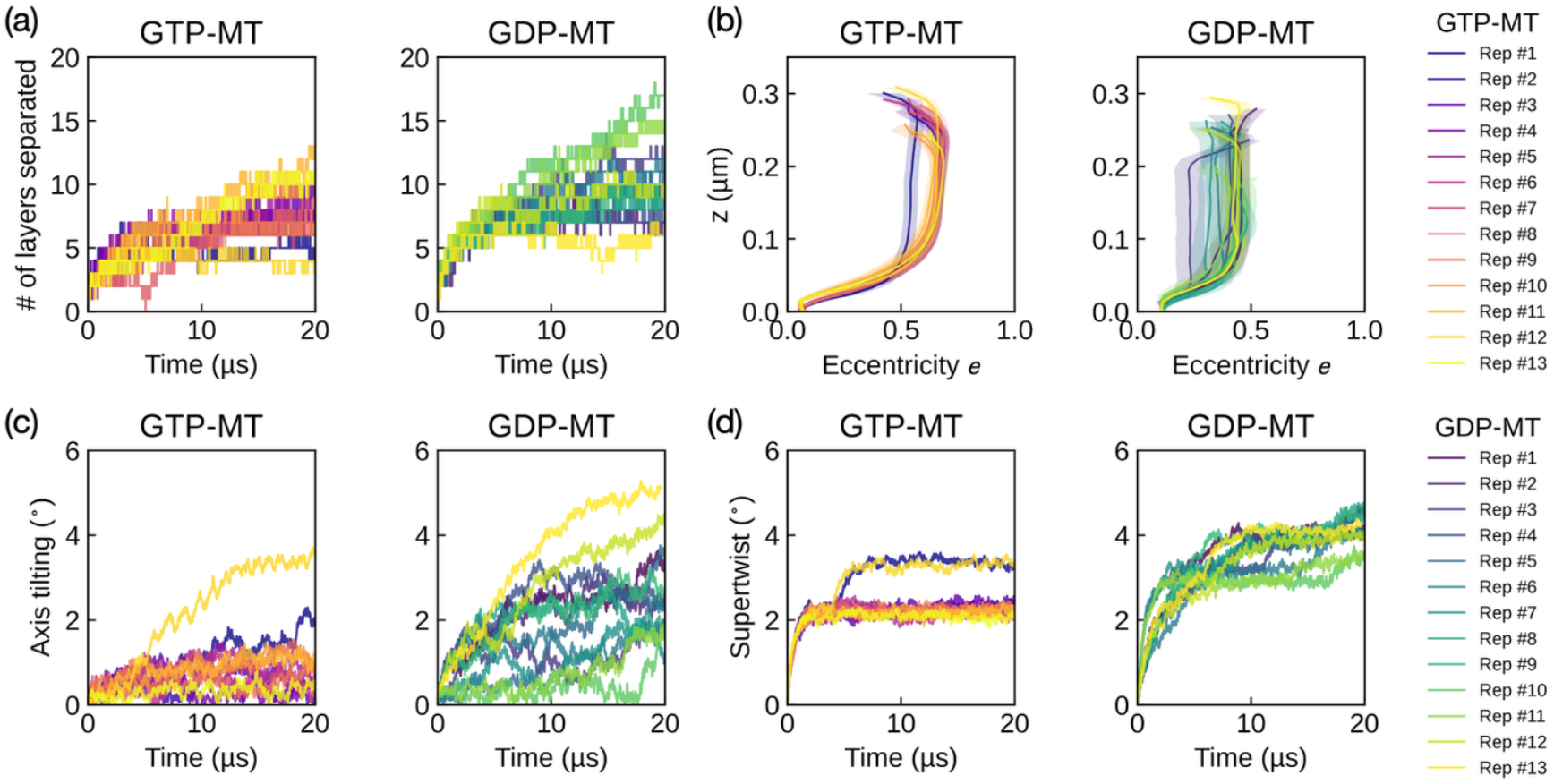
Analysis of CG MD simulation results on 40-layer 14-PF MT models. Replicas 1-9 are with 2 layers of bottom restraints; replicas 10-13 are replicas 1-4 shown in **Fig. 6**. For 13-PF statistics, see Figure S1.14. (a) Time evolution of number of layers that splits to form clusters (“peels”) in simulations, ***n***_***s***_(***t***). (b) Layer eccentricity along protofilament ***z*** (axial) coordinates; error bars calculated for last 5 µs of each replica trajectory. Plotted from bottommost unrestrained layer to the peeling layer. (c) MT axis tilting angle as a function of time. The MT axis tilting angle is defined as the angle between the MT axis direction and the ***z*** axis (initial MT axis) direction. The MT axis direction is fitted from the COMs of layers, from the bottommost unrestrained layer to the uppermost unpeeled layer. (d) MT supertwist angle as a function of time. The supertwist angle is defined as the pointing direction of each protofilament and the MT axis direction calculated in (c). The PF direction is fitted from the COMs of tubulin heterodimers, from the bottommost unrestrained layer to the uppermost unpeeled layer.

#### (3) Lattice tilting

It is also in these two replicas that we observed bent curvature along the MT lattice that caused a tilt away from the z-axis (**Fig. 7c**). We defined the axis tilting angle as the angle between the MT axis direction and the z axis (initial MT axis) direction. The MT axis direction is fitted from the COMs of layers, from the bottommost unrestrained layer to the uppermost unpeeled layer. Such collective tilting may be another mechanism to release MT lattice strain and may be responsible for stabilizing the peeled MT tip at limited layers. On average, GDP-MT exhibited more pronounced lattice tilting behavior, hinting at larger lattice strain to release. This trend also applies to 13-PF results in **Fig. S1.14**.

#### (4) Protofilament in-lattice supertwist

There is in-lattice leftward supertwist of PFs in both GTP-MT and GDP-MT (**Fig. 7d**). We calculated the supertwist angle as the average angle of each protofilament direction with the MT axis direction. Such in-lattice torsion is not observed in the 8-layer model due to the existence of positional restraints, and the leftward direction of supertwist is in accordance with torsion of PF clusters in both AA and CG MD simulations. This observation also aligns with the Moiré patterns observed in experiments (47). We also observed significantly larger supertwist in GDP-MT than GTP-MT. For GTP-MT, a small number of replicas (reps 1,12 of 14-PF, reps 1, 2, 7 of 13-PF) also exhibited an increase into GDP-like supertwist while having fewer layers peeled than other GTP-MT replicas (**Fig. 7d, Fig. S1.12-14**). We hypothesize that such in-lattice torsion is another mechanism for the MT lattice to release lattice strain in the lower lattice layers that cannot undergo lateral dissociation due to neighboring interactions, and such relief of torsion slows down or even counteracts further peeling at the MT tip.

#### (5) Elliptical deformation in the GTP cap

The *z*-axis projections of the unpeeled lattice layers can slightly depart from a circular shape to an elliptical shape, especially in GTP-MT (**Fig. 7b**). For each heterodimer layer *j* beneath the peeling layer, we projected the layer along the local MT axis. The local MT axis direction at layer *j* is determined by fitting a line through the COMs of the neighboring heterodimer layers (*j* − 2) to (*j* + 2). The projected COMs is fitted into an ellipse via least-squares optimization, and the is determined. We can see that GTP-MT undergoes more elliptical deformation than GDP-MT, thus favoring flatter, more rigid and thus more stable PF clusters, supporting the conclusions in (32).

These results facilitate explanation of the origin and the role of the GTP cap at MT plus ends. Despite its lattice elongation (15), the GTP cap is structurally more rigid. The stronger lateral interaction in the GTP cap forms shorter and less PF clusters on average, which stabilizes the bulk lattice by limiting destabilizing lateral dissociation events, limiting solvent exposure of tubulins. The elliptical deformation in the GTP cap adds to the structural rigidity of that cap, since it shows that GTP-tubulins may favor a flatter contact geometry than GDP-tubulins in both the lattice and the PF cluster. This favors the stability of such PF clusters, limits their dissociation, and thus favoring growth. This also adds a barrier to the hydrolysis event, as the newly hydrolyzed tubulins will also need to overcome a slight in-plane relaxation from this more elliptical shape. However, for MTs to exhibit their flexibility and mobility and to facilitate intracellular transport, a cylindrical shape and the greater flexibility of the GDP-MTs is more favored for the bulk MT lattice than the GTP-MT.

### All-atom mechanism for PF cluster formation and stabilization

After our thorough CG modeling, we now aim to understand the atomistic origins of the formation and stabilization of the PF clusters. How do the residue contacts at these interfaces, at a faster timescale, affect the collective dynamics in a slower timescale? What interactions are the most relevant in this process? Through our analysis, we identified key residue contacts, uncovered a separation of timescales in the relaxation of longitudinal and lateral interfaces, and proposed a mechanism for PF cluster formation and stabilization.

We first analyzed the lateral interface in much detail, which should be responsible for the nucleotide difference in the lattice model. We first analyzed the average number of hydrogen bonds (H-bonds) in each undissociated lateral pair, in different segments of the trajectory (**Figs. S2.1-S2.3**). Surprisingly, there was no significant difference between the two nucleotide states (*p* = 0.638), indicating the lateral interactions, when solely identified as H-bonds, are very similar across nucleotide state, and the all-atom sampling noise across different heterodimers in the same lattice is more significant than the average difference between the two nucleotide states. We then investigated the most frequent H-bond pairs in the *β*-*β* and *α*-*α* lateral interactions (**Fig. 8a**) and calculated their frequency correlation averaged among different lateral pairs (**Figs. S2.4, S2.5**). While still no significant difference for GTP-MT and GDP-MT was found, our analysis revealed important structural insight in the process of the bending relaxation of PF clusters. First, we noticed that the strongest H-bonds in the lateral interface form between H9/H10 helices in PF-*n* and the H3 helix in PF-(*n*+1). Specifically, in *β*-tubulin, there is a H-bond “quadrupole” among *β*:Asp295(*n*), *β*:Lys297(*n*), *β*:Asp118(*n*+1) and *β*:Lys122(*n*+1) (**Figs. 8b, S2.8a**), characterized by their positive probability correlation (**Fig. S2.4**), indicating large cooperativity in their formation. In α-tubulin, while there is no such “quadrupole”, the *α*:Glu297(*n*)-*α*:Lys124(*n*+1) H-bond has a larger frequency of formation (**Fig. S2.5**) compared with *β*-tubulins (**Fig. S2.4**), indicating comparable lateral interaction strengths. These H-bond residues are also capable of forming salt bridges to enhance their interaction. To our surprise, the H-bonds formed between the M-loop and H1/S2-loop and H2/S3-loops are weaker than those between H9/H10 helices and H3 helices in *β*-tubulin, contrary to the structural characterizations in (9), and this result is in agreement with insight from our CG forcefield (**Fig. S1.4**). In *α*-tubulin, *α*:Lys280(*n*) and *α*:Glu284(*n*) in M-loop form stronger H-bonds with the H2-S3 loop than those in *β*-tubulin. Their simultaneous presence requires the sidechains of Lys280 and Glu284 to be in the same direction (**Fig. S2.8b**), favoring an *α*-helical secondary structure of the M-loop, as shown by DSSP analysis (**Fig. S2.8d**).

**FIGURE 8.**
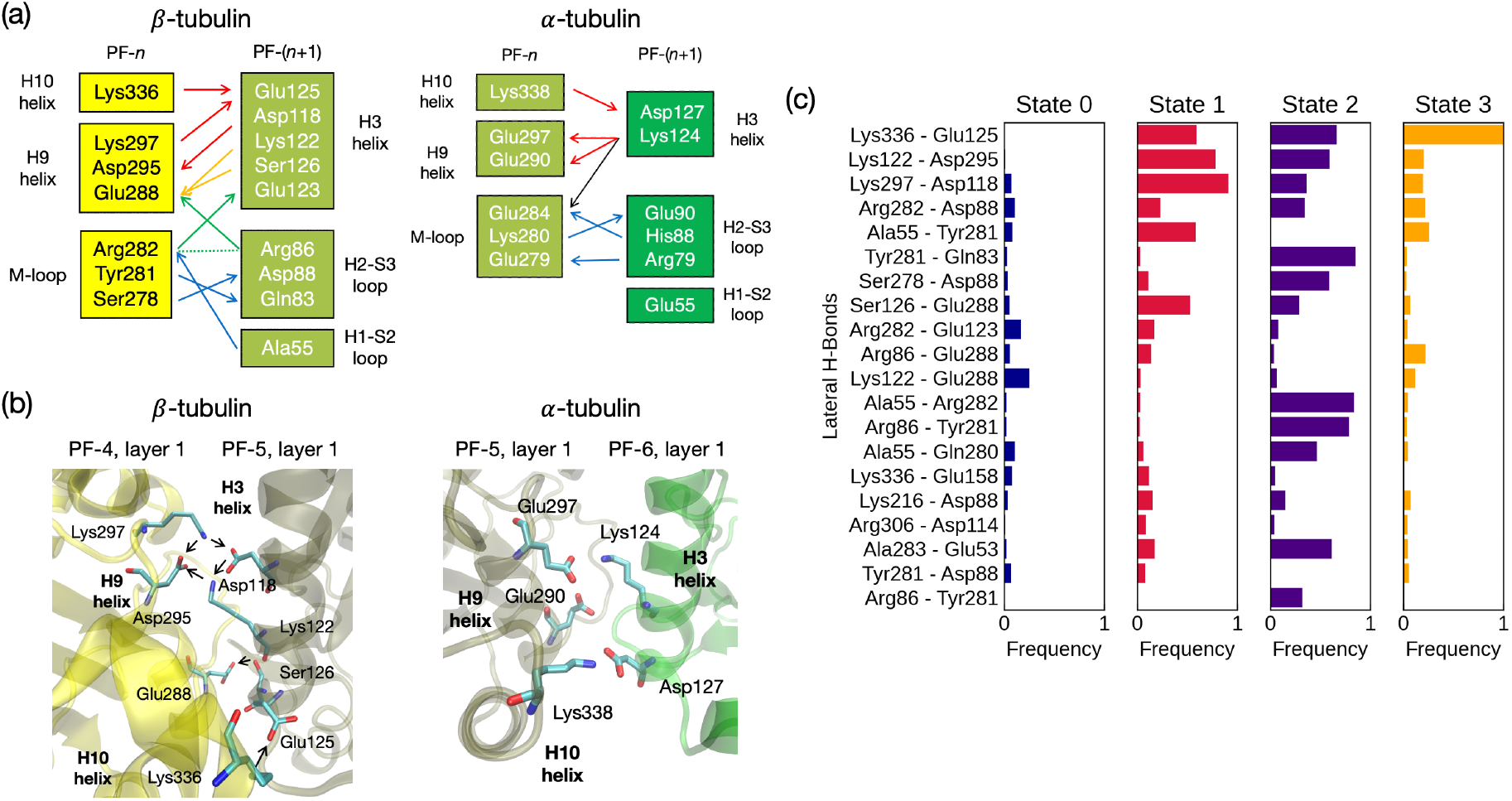
All-atom analysis of homolytic lateral interaction. (a) Dominant H-bond interaction networks of β-tubulin and α-tubulin. Different colors correspond to separate illustrations of interaction in the Supporting Information. Arg-Arg stacking interaction in β-tubulins is denoted by a dashed line. (b) Illustration of H9/H10-H3 interaction in β- and α-tubulins, with examples from PF-4 & PF-5 and PF-5 & PF-6 respectively. (c) Dominant H-bond interaction modes in β-tubulin, characterized by PCA/K-Means analysis.

Because *β*-*β* lateral interface had more diverse H-bond pairs than *α*-*α* interface, we used Principal Component Analysis (PCA) and K-Means Clustering to identify the most dominant binding modes in the *β*-*β* lateral interface, in the linear space spanned by the top 20 most frequent H-bonds, as described in Methods. We identified 4 as the optimal cluster number (**Fig. S2.9**) and found that each state represents different features in lateral interaction. Across the 8-layer MT lattice and along the 4 µs trajectory, each *β*-tubulin can transition from these 4 different lateral interaction modes stochastically (**Figs. S2.10-S2.13**). State 0 corresponds to a dissociated state; state 3 corresponds to a near-dissociated state with only the H10-H3 H-bond pertaining; states 1 and 2 represent characteristic “PF cluster style” and “lattice style” interactions respectively, because: (a) during the 4 µs simulation, state 1 became more frequent while state 2 became less frequent among all lateral pairs (**Figs. S2.11, S2.12**); (b) state 2 is characterized by strong contribution by interaction of M-loop with the H1/S2 and H2/S3 loops (e.g., H-bond pairs of *β*:Arg282(*n*+1)-*β*:Ala55 and *β*:Tyr281(*n*+1)-*β*:Arg86; see **Fig. 8c**), which agrees with the lateral interaction in the MT lattice state characterized by ref. (9); conversely, state 1 lacks such interactions but has larger contributions from H9-H3 interaction, characterized by higher frequencies in “quadrupole” H-bonds and the *β*:Glu288(*n*)-*β*:Ser126(*n*+1) H-bond (**Figs. 8c, S2.4**), serving to “lock” the opposite sides of the helices. It is also noteworthy that the latter H-bond increases in frequency rather than decreases during simulation, indicating its possible stabilizing effect of the lateral interface in MT clusters.

The relatively stronger interaction between H9 and H3 helices prompted us to analyze the torsional angle and distances between the two helices. We found that, in the first 0.5 µs, the bimodal helical distance distribution in β-tubulin has already shifted (**Fig. S2.15**), favoring a closer contact. The crossing angle between H9 and H3 helices flattened in a longer timescale in both α- (**Fig. S2.14**) and β-tubulins (**Fig. 9**). These results together demonstrate that the continued outward bending relaxation of PF clusters may be a result from structural relaxation in the lateral interface. However, it is noteworthy that the H9-H3 contact between different lateral pairs is rather stochastic than uniform, and the characteristic timescale for lateral relaxation is obviously longer than that of the initial cleavages that drove the formation of PF clusters, which lies within 0.1 µs (**Fig. S2.19**). Only the initial jump of longitudinal inter-dimer (**Fig. S2.20**) and intra-dimer (**Fig. 10b**) distances (both of which we abbreviate as “longitudinal jump”) matches within this timescale.

**FIGURE 9.**
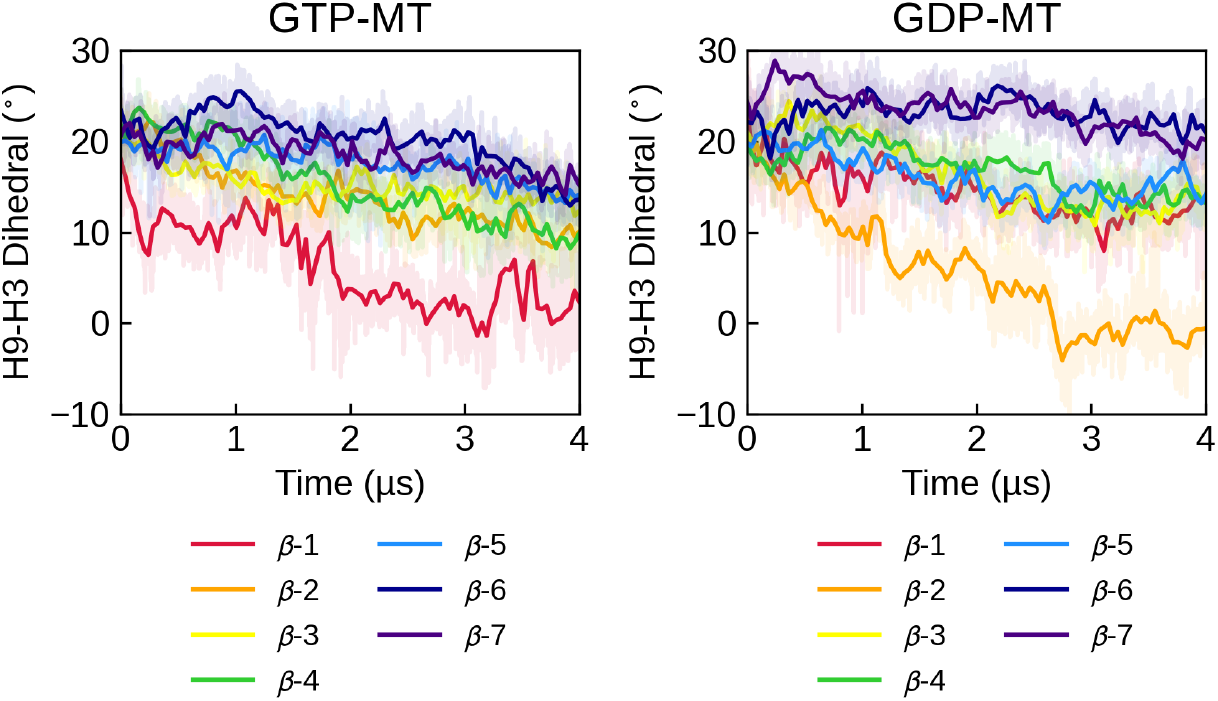
Time evolution of the mean H9-H3 dihedral angle in β-tubulin (see Fig. S2.14 for α-tubulin) in each heterodimer layer.

**FIGURE 10.**
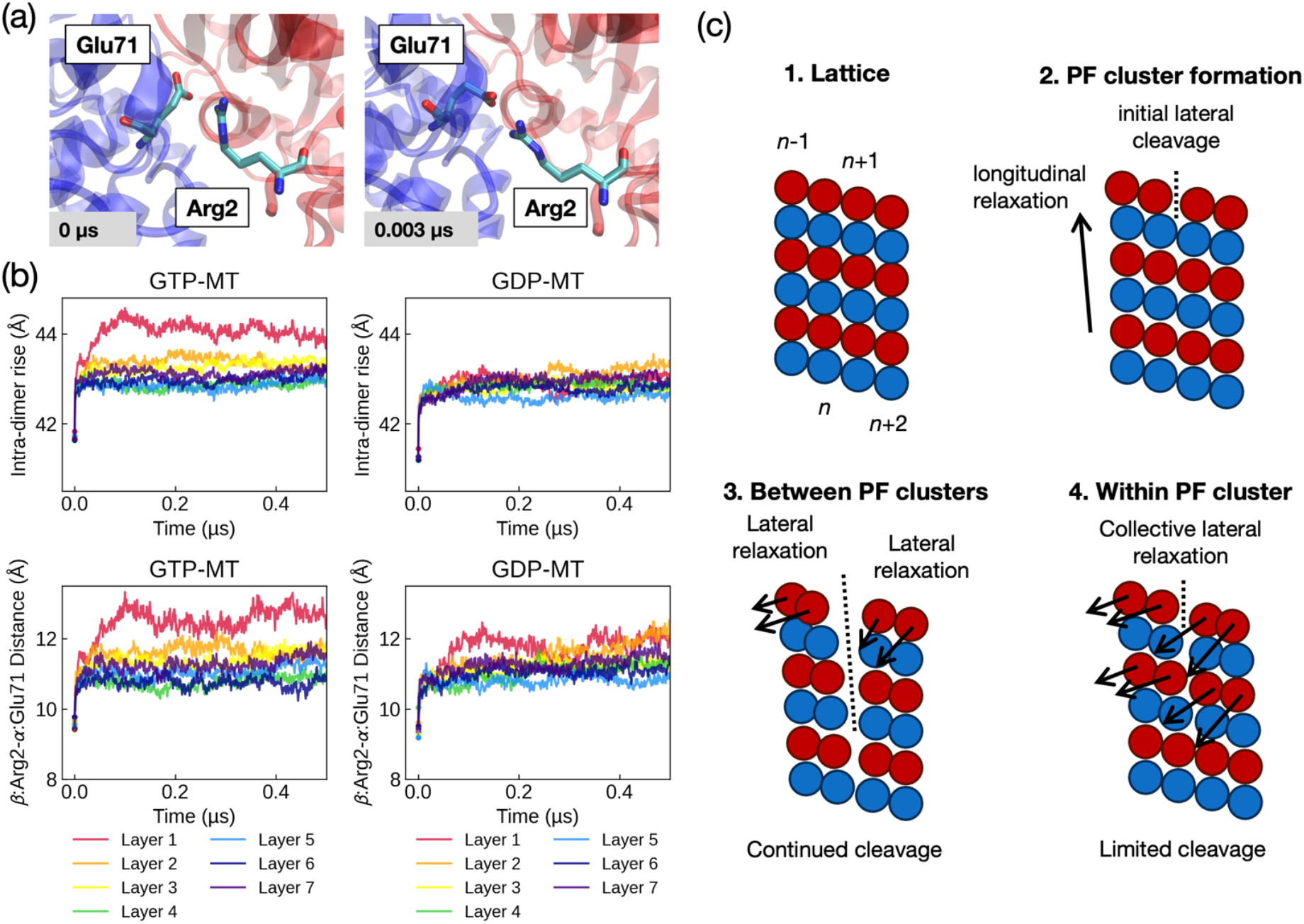
All-atom MD analysis of longitudinal interaction and mechanism for PF cluster formation. (a) Illustration of the relaxation of intra-dimer β:Arg2-α:Glu71 H-bond. Plotted for Layer 7 of PF1. B-tubulin (red) moves further away from α-tubulin, especially obvious in N-terminus. (b) Time evolution of average intra-dimer COM distance in each heterodimer layer. (see Fig. S2.20 for inter-dimer distance), and the average Cα-Cα distance in the β:Arg2-α:Glu71 H-bond pair, synchronous to the intra-dimer distance jump. (c) Illustrated mechanism for PF cluster formation and stabilization. The radial component of the longitudinal jump promotes initial lateral cleavage, and subsequent lateral relaxation promotes continued cleavage between PF clusters, stabilization within PF clusters, and collective outward bending motion of PF clusters. Subsequent openings at the tip (**Fig. 3**) are limited by lateral relaxation in lower layers.

With this in mind, we analyzed the longitudinal interfaces (**Figs. S2.6, S2.7**) and examined whether any of the H-bond pairs facilitate the longitudinal jump. For the intra-dimer H-bonds, we noticed that the *β*:Arg2-*α*:Glu71 H-bond (**Fig. 10a**) formed in the first 3 ns via sidechain relaxation, and the stabilization of such H-bond requires sidechain stretching and orientation matching. This is manifest in the jump in Cα-Cα distance of this H-bond pair (**Fig. 10b**), which is surprisingly uniform across the lattice and synchronous to the longitudinal jump (**Fig. 10b**) in each heterodimer layer both in trend and in magnitude (both increased by ~2 Å). In the inter-dimer interface, there is also a similar H-bond pair α:Arg2-β:Glu69 (**Fig. S2.7**). A similar pattern is also observed for the β:Lys324-α:Glu207 H-bond (**Fig. S2.22**), where the formation of such H-bond also require sidechain stretching. While the longitudinal jump can also be caused by thermal fluctuation and is not necessarily a result of H-bond relaxation, the formation of such strong H-bonds undoubtedly reinforces the initial structural perturbation. Such structural perturbation happens not only in the z-direction, but in the r-direction as well (**Fig. S2.23**), and initiates the earliest lateral cleavage from this outward bending tendency at the MT plus end.

Therefore, we can sketch a complete physical picture in **Fig. 10c** for the formation and stabilization of PF clusters at the all-atom resolution. When the system relaxes from its low temperature starting structure, increased thermal fluctuations enable an initial expansion of the microtubule lattice. This structural transition is then stabilized by the formation of strong longitudinal hydrogen bonds that favor a more expanded configuration. This also creates the earliest lateral cleavages at the PF breaking points, which in turn provides opportunity for closer H9/H10-H3 contacts in the lateral interface within the PF clusters. The closer and more aligned H9-H3 contact sustains more outward and leftward bending in PF-(*n*+1) than PF-*n*, when *n* is not a breaking point. When it is a breaking point, PF-(*n*+1) and PF-*n* experience different torsional movement due to lateral relaxation in the neighboring PF clusters, so PF-(*n*+1) and PF-*n* continues to split apart. Such continued splitting in turn reinforces the lateral interaction within PF clusters, so they remain stable. Even if partial lateral dissociation is observed at the top of the MT tips in the all-atom simulations, they remain limited due to lateral relaxation in lower layers, so that the MT tip configurations, formed in the first 0.5 µs, remain stable up to 4 µs. While the seam line is the most constant breaking point in terms of thermodynamics and statistics (as in **Fig. 5c**), its lateral cleavage is neither the first nor the fastest in terms of dynamics among all possible breaking points (as shown in **Fig. S2.19**).

These results, when combined with CG MD simulations, reveal that the intrinsic thermodynamic instability of the MT lattice drives the formation of stable PF clusters at the MT plus end. From the CG perspective, such an instability is manifest in the learned curvature directly from PF cluster layers. Under equilibrium sampling of the CG forcefield, the system tries to relax anywhere in the lattice, so we see the PF clusters in the 8-layer model remain stable and clustered, while the 40-layer model can relax by: (a) continuing to splay out; (b) in-lattice twisting; or (c) developing curvature of MT axis spontaneously, thus enabling MT mobility and flexibility within the cell. The latter two means of relaxation can therefore – along with dynamical friction and entropic penalty from increased solvent exposure – stall the outward peeling of MT plus-end tips and stabilize the lattice layers. From the AA perspective, the rapid transition from lattice interaction modes to the interaction modes in PF clusters indicates the instability of lattice-style interaction modes at the simulated temperature, and such a relaxation process eventually drives the continued outward bending and stabilization of PF clusters. The local equilibrium sampling approximation encoded in the bottom-up CG methodology is also justified by our all-atom analysis that the interaction interfaces in PF clusters are indeed more relaxed. Although the longitudinal and lateral H-Bond interactions are similar between GTP-MTs and GDP-MTs, this does not contradict the weaker lateral interaction that resulted from the CG modeling, because: (a) other types of interactions and geometric considerations can be at play, and (b) the lateral interaction modes are stochastic across the lattice, resulting in large noise in the all-atom MD data.

## CONCLUSIONS

Microtubule catastrophe is fundamentally driven by a thermodynamic force: the inherent instability of the MT lattice. In this work, we have provided a comprehensive multiscale study of microtubule tip relaxation dynamics and PF cluster formation through a bottom-up CG model for tubulins, constructed from 4 µs all-atom MD data for an 8-layer MT lattice. To our knowledge, this work is the first to conduct extensive CG modeling of long MT systems with a bottom-up CG model in 20-site-per-monomer resolution, higher than recent CG models of MTs (29, 32, 48), the first to delve into atomistic structural details in the room-temperature MD data of an entire MT lattice, and the first to thoroughly examine the origin and fate of the via direct AA and CG simulation.

Our simulations and analysis reveal that PF clusters are likely to be stable, long-lived intermediates in both MT growth and decay. The straight MT lattice is not stable and relaxes to the PF clusters at the MT-tip, indicating a spontaneous breaking of z-axis helical symmetry. In the 8-layer models, The PF clusters remain stable up to 40 µs of CG simulation time and the clustered PFs exhibits collective bending and torsional motion. In the 40-layer models, such lattice instability becomes more pronounced, and PF clusters continue to peel and bend out up to 20 µs of CG simulation time. Their formation can be limited by other means of strain relaxation by lattice curvature and supertwist. The equilibrium configurations of PF clusters are not deterministic. Their grouping may vary, although some configurations are more probable. Though not frequent in the simulations, rearrangement of PF cluster configurations and lateral suturing/peeling fluctuations are also observed, further demonstrating the conformational flexibility of these PF clusters.

The formation of PF clusters is a hierarchically coordinated process by relaxations in the longitudinal and lateral interfaces that involves different timescales. The longitudinal interface forms stronger H-bonds in general, relaxes faster, and is more consistent across different tubulins inside the lattice. The lateral interface forms weaker H-bonds but exhibits much larger variability and multimodal behavior, so it relaxes more slowly. On the nanosecond timescale, relaxation of longitudinal H-bond pairs promotes lattice elongation, which initiates the first few lateral dissociations at the plus-end tip. Then, on the microsecond timescale, lateral relaxation stabilizes the PF tips by switching of H-bond interaction modes, especially between H9/H10 and H3 helices (49). The resulting alignment of H9 and H3 helix orientations acts as a major structural difference between the lateral interfaces inside PF clusters and in the MT lattice.

The fate of PF clusters, as a part of a physiological-temperature MT plus-end tip, is with the microtubule itself, both governed by its nucleotide state and the resulting difference in lateral interaction. By directly tuning the lateral interaction strengths, we see a clear trend that the number of PF clusters increases and bending relaxation accelerates when the lateral interaction weakens. As shown in the CG modeling, GDP-MTs have weaker lateral interactions, leading to faster bending relaxation and a higher propensity for lateral cleavage. More PF clusters, a lateral event, eventually enable easier depolymerization in the longitudinal direction, by preemptively overcoming the lateral dissociation barrier and solvent rearrangement barriers, both required for a tubulin to fully detach from the tip. GTP-MTs, on the other hand, have stronger lateral interactions in the PF cluster, which can in turn hold more PFs together in a somewhat flatter conformation. This favors growth by preemptively overcoming the solvation rearrangement barrier during the tubulin-tubulin, tubulin-PF, and PF-PF association events. This physical picture concurs with the “conformational selection” mechanism proposed by ref. (32) if we define “conformational states” with the number of laterally dissociated PF pairs (i.e., the number of PF clusters in this article). That is, conformation states with more split PF pairs are more structurally volatile, more exposed to solvent, and more prone to undergo longitudinal dissociation without the neighboring interactions. GDP-MTs have a larger propensity to occupy these states, thus making shortening events more possible. The reversibility of PF clustering and lateral dissociation events in our 40-layer simulations also supports this “conformational selection” mechanism.

Our results also highlight the existence and effect of PF supertwist in MT simulations. To our best knowledge, all MD simulations have used cryo-EM structures of MT lattice patches to construct a MT lattice with translational symmetry in the z-axis, but both our simulations and the previous results on a 6-layer MT tip by Igaev and Grubmüller (31) have exhibited the formation of stable PF clusters and rupture of the MT plus end from the start of the all-atom MD simulations. It is therefore likely that translational symmetry in the z-axis is broken under higher temperature in both GTP- and GDP-MT lattices and, therefore, starting the simulation from a lattice directly constructed from cryo-EM structure may suffer from greater lattice instability and exhibit less realistic relaxation dynamics. Recent work utilized a Martini CG model for the MT lattice and the authors proposed that a longitudinal supertwist of around 2 degrees per dimer must be in place to stabilize the lattice from rupture (50). However, we note that the Martini model fails to correctly capture the enthalpy-entropy decomposition of pair correlations so these results may not be completely reliable (51, 52). This Martini MT model also suffers unphysical artifacts such as in-lattice ruptures and must be combined with a manually tuned Elastic Network Model (ENM) to stabilize its MT lattice structure. It will be a goal of our future work to determine the extent of lattice twist in an all-atom MD model and whether the lack of such a lattice twist in the initial modeling overemphasized the formation of PF clusters.

We conclude this paper by noting some limitations of our current CG model. We approximated the clustered layers as locally equilibrated in 3-4 µs of AA MD, so that we could infer a local potential of mean force from them and model the MT relaxation process. However, full global relaxation remains elusive for a reference AA trajectory in a system this large. The resolution of the CG model is also relatively high (a larger number of CG sites or beads), so we are currently unable to simulate long MTs on a long timescale to observe dissociation or association. It will also be a target for our future work to simulate mixed-state lattices (e.g., a GDP-MT lattice with a GTP cap) and to develop Ultra-Coarse-Grained (UCG) models (53–55) that can simulate dynamic nucleotide state transitions (56, 57) within the lattice environment as the simulation trajectory evolves.

## DATA AND CODE AVAILABILITY

Sample LAMMPS input files and trajectories for CG simulations are provided in the online Zenodo server https://doi.org/10.5281/zenodo.16995325. Other scripts and data are available upon request to the authors.

## SUPPORTING MATERIAL

Supporting Information can be found online, containing one supporting document (Table S1.1, Figures S1.1-S1.7, Figures S2.1-2.23) and supporting movies.

## AUTHOR CONTRIBUTIONS

W.X., J.W., T.B., and G.A.V. conceptualized the research. W.X. performed the CG model development, CG simulations, and AA MD analysis, and wrote the initial draft of the manuscript. All the authors have analyzed the data and contributed to the editing of the final version of the manuscript.

## ACKNOWLEDGMENTS

This work was supported by the US Department of Energy, Office of Science, Basic Energy Sciences, under award DE-SC0023318. We gratefully acknowledge computing time provided by the University of Chicago Research Computing Center (https://rcc.uchicago.edu), the University of Chicago high-performance GPU-based cyberinfrastructure supported by the National Science Foundation under grant no. DMR-1828629, and Frontera at the Texas Advanced Computing Center (TACC) at The University of Texas at Austin (http://www.tacc.utexas.edu) funded by the NSF (OAC-1818253), and the NIH-funded Beagle3 HPC cluster (Award Number S10OD028655).

## DECLARATION OF INTEREST

There is no conflict of interest to declare.

## Notes

### Competing Interest Statement

The authors have declared no competing interest.

### Summary of Updates

New coarse-grained simulations conducted for 13-protofilament models as requested by the reviewers; Updated abstract, introduction, results and discussion, conclusions, in response to reviewers

