## Supplementary Information for "Lattice Instability Drives Formation of Protofilament Clusters at the Microtubule Plus End Tips"

#### Contents for All Supporting Information

**Supporting Document (This Document)**

Section 1: Coarse-Graining (Table S1.1, Figures S1.1-S1.7)
Section 2: All-Atom (Figures S2.1-2.23)

**Supporting Movies (movies.zip)**

Movies S1.1 – S1.10: CG MD continuations on the 4 μs AA configuration
Movies S2.1 – S2.8: CG MD trajectories for the 40-layer model

**Data And Code Availability**

Sample LAMMPS input files and trajectories for CG simulations are provided in the online Zenodo server https://doi.org/10.5281/zenodo.16995325. Other scripts and data are available upon request to the authors.

### Supporting Document

#### Section 1: Coarse-Grained Modeling

**Table S1.1** Coarse-grained (CG) mapping for alpha- and beta-tubulins from KMC-CG.

| Beta-tubulin | | Alpha-tubulin | |
|---|---|---|---|
| **CG site ID** | **Residues** | **CG site ID** | **Residues** |
| 1 | 327-350 | 21 | 34-36, 46-48 |
| 2 | 243-248 | 22 | 389-400, 418-427 |
| 3 | 428-429 | 23 | 37-45 |
| 4 | 137-140, 166-173, 180-202, 302-304 | 24 | 285-293, 324-338 |
| 5 | 94-95 | 25 | 95-118, 144-153 |
| 6 | 249-268, 305-314, 368-374 | 26 | 138-143, 168-205, 387-388 |
| 7 | 401-408 | 27 | 401-417 |
| 8 | 21-62, 82-84 | 28 | 259-269, 310-316, 339-352, 378-386, 428-438 |
| 9 | 390-400 | 29 | 241-248 |
| 10 | 96-98 | 30 | 279-284 |
| 11 | 72-76 | 31 | 236-240, 249-258, 317-323, 353-361, 371-377 |
| 12 | 375-389, 409-427 | 32 | 1-7, 119-137, 165-167 |
| 13 | 276-280 | 33 | 154-164 |
| 14 | 1-5, 117-134, 155-165 | 34 | 8-33, 49-52, 64-94 |
| 15 | 99-116, 141-154 | 35 | 206-235, 270-278, 294-304 |
| 16 | 174-179 | 36 | 439-441 |
| 17 | 234-242, 269-272, 281-294, 315-326, 351-367 | 37 | 53-56, 60-63 |
| 18 | 6-20, 63-71, 85-93, 135-136 | 38 | 57-59 |
| 19 | 203-233, 273-275, 295-301 | 39 | 362-370 |
| 20 | 77-81 | 40 | 305-309 |

**Figure S1.1:** KMC-CG results for 14 CG sites. Note that the M-loop is not individually defined in both alpha- and beta-tubulins.

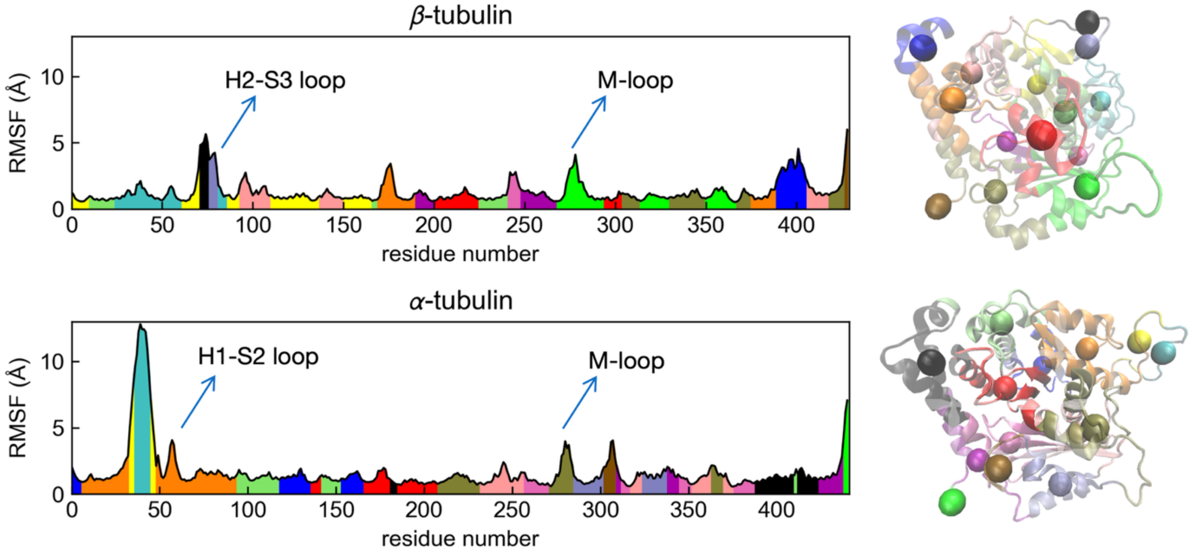


**Figure S1.2:** KMC-CG results for 18 CG sites. Note that the M-loop is not individually defined in alpha-tubulin.

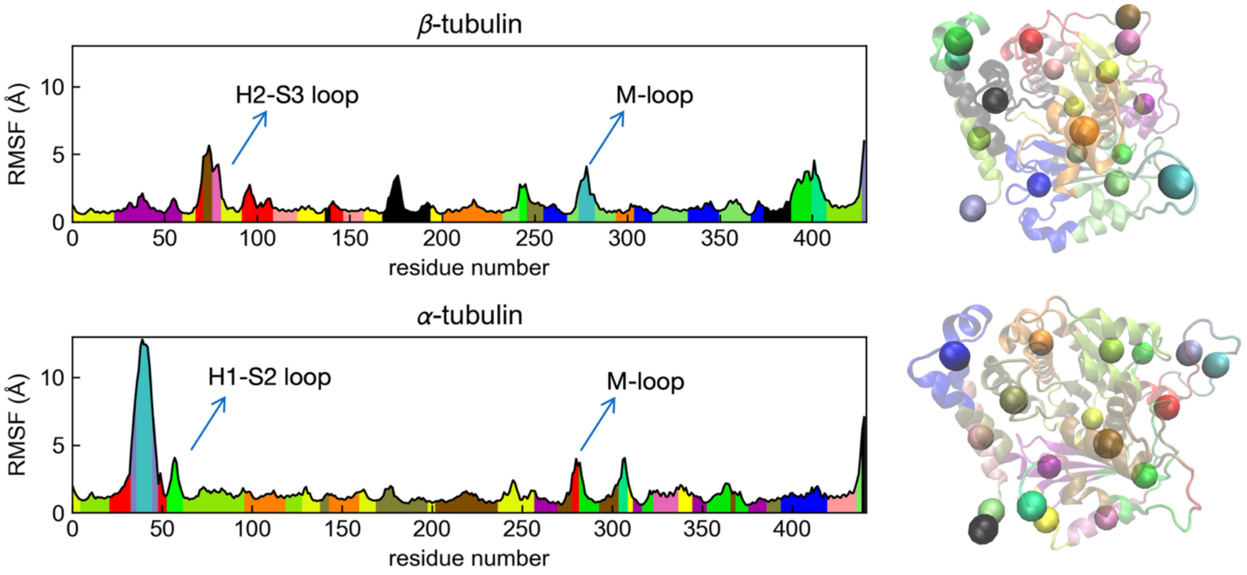


**Figure S1.3:** Comparison of longitudinal interaction intensities in the CG forcefield.

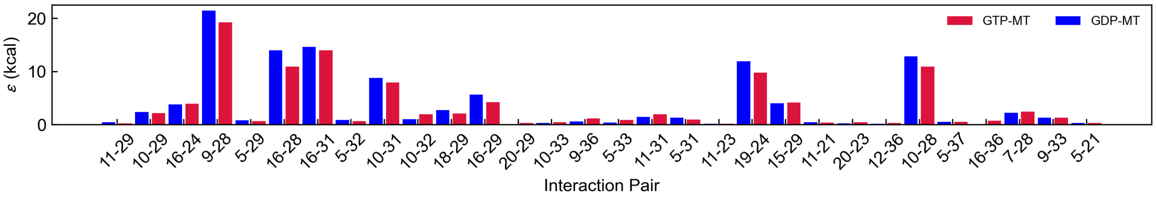

**Figure S1.4:** Comparison of lateral interaction intensities in the CG forcefield.

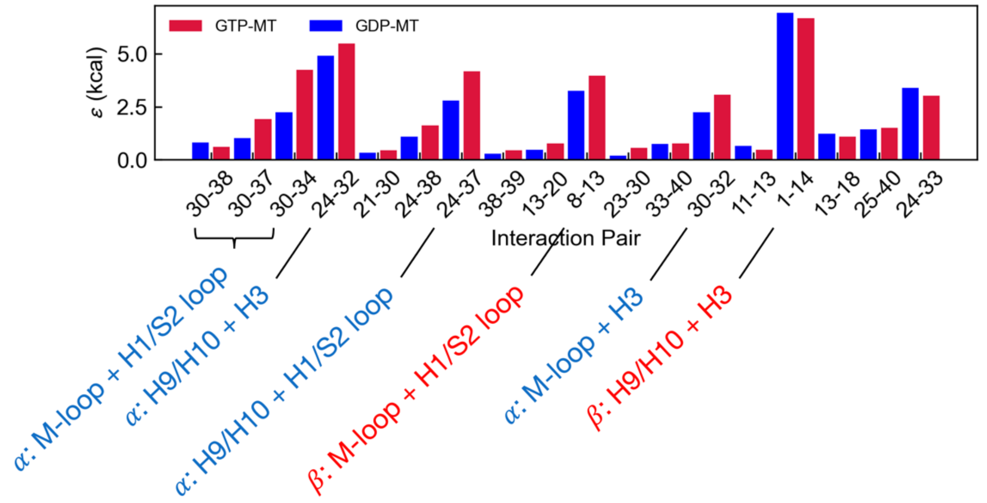


**Figure S1.5:** Averaged bending curves for adjusted damping factors.

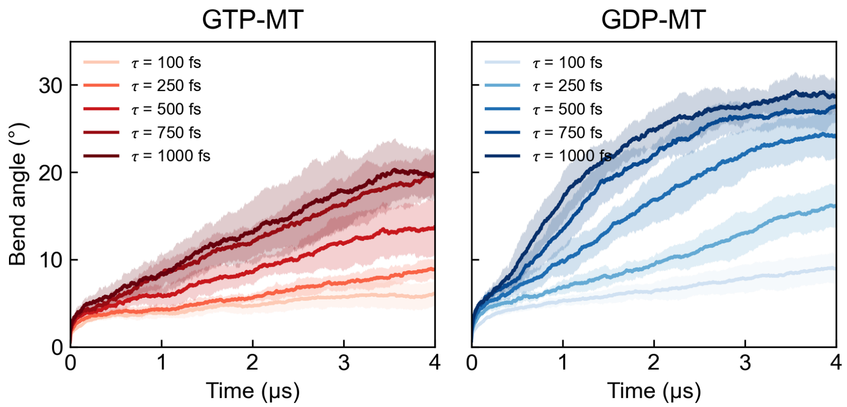


**Figure S1.6:** Averaged bending curves for scaled lateral interaction.

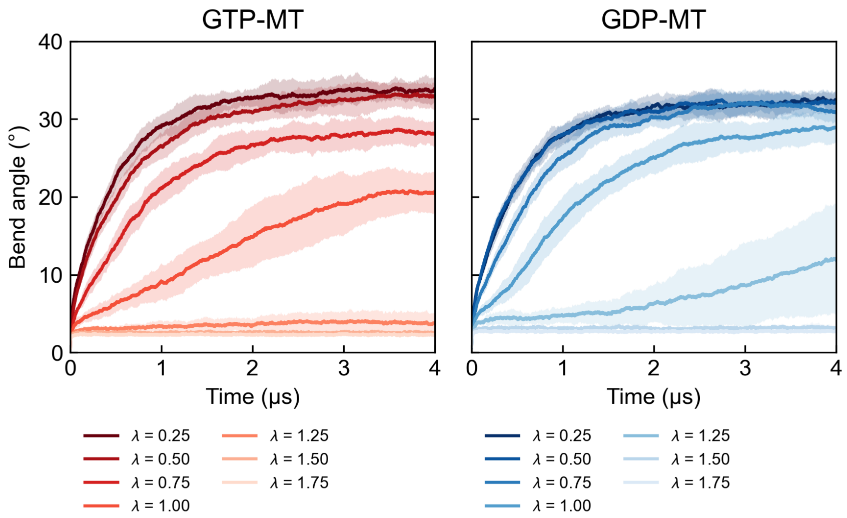

**Figure S1.7:** Most frequent PF cluster configurations from default forcefield parameters.

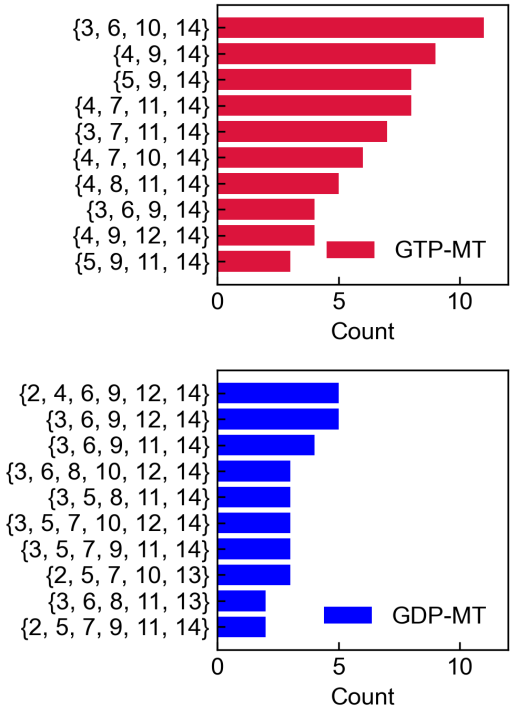


**Figure S1.8:** Evolution of intra-dimer distances in the MT system, all-atom compared with CG model.

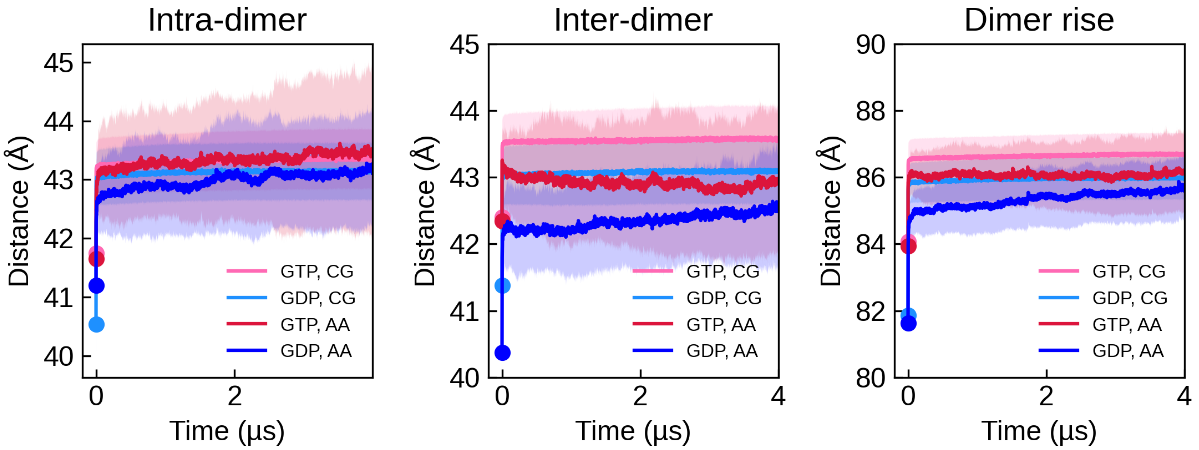

**Figure S1.9:** Evolution of intra-dimer distances in the MT system, all-atom compared with CG model.

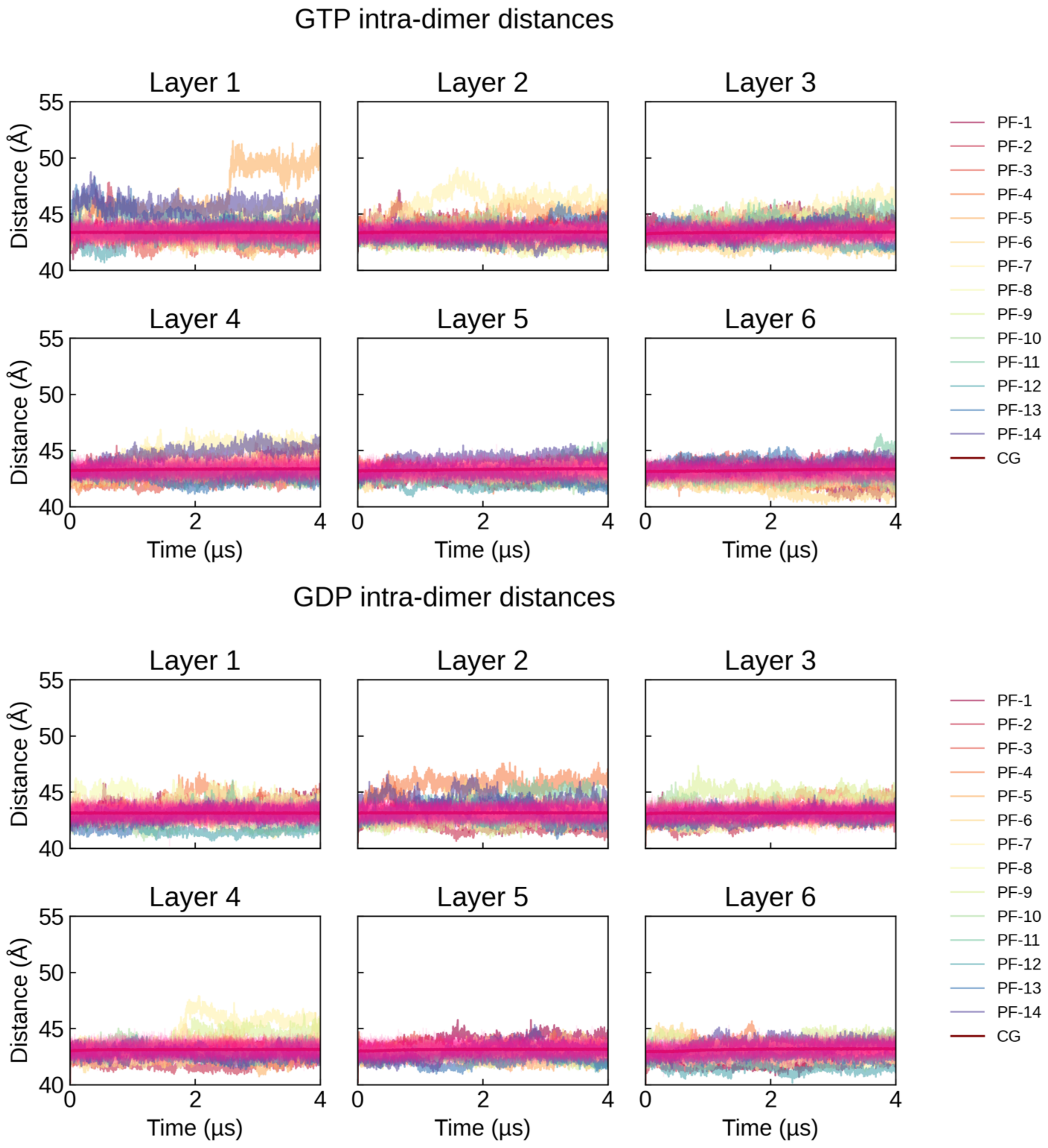

**Figure S1.10:** Evolution of inter-dimer distances in the MT system, all-atom compared with CG model.

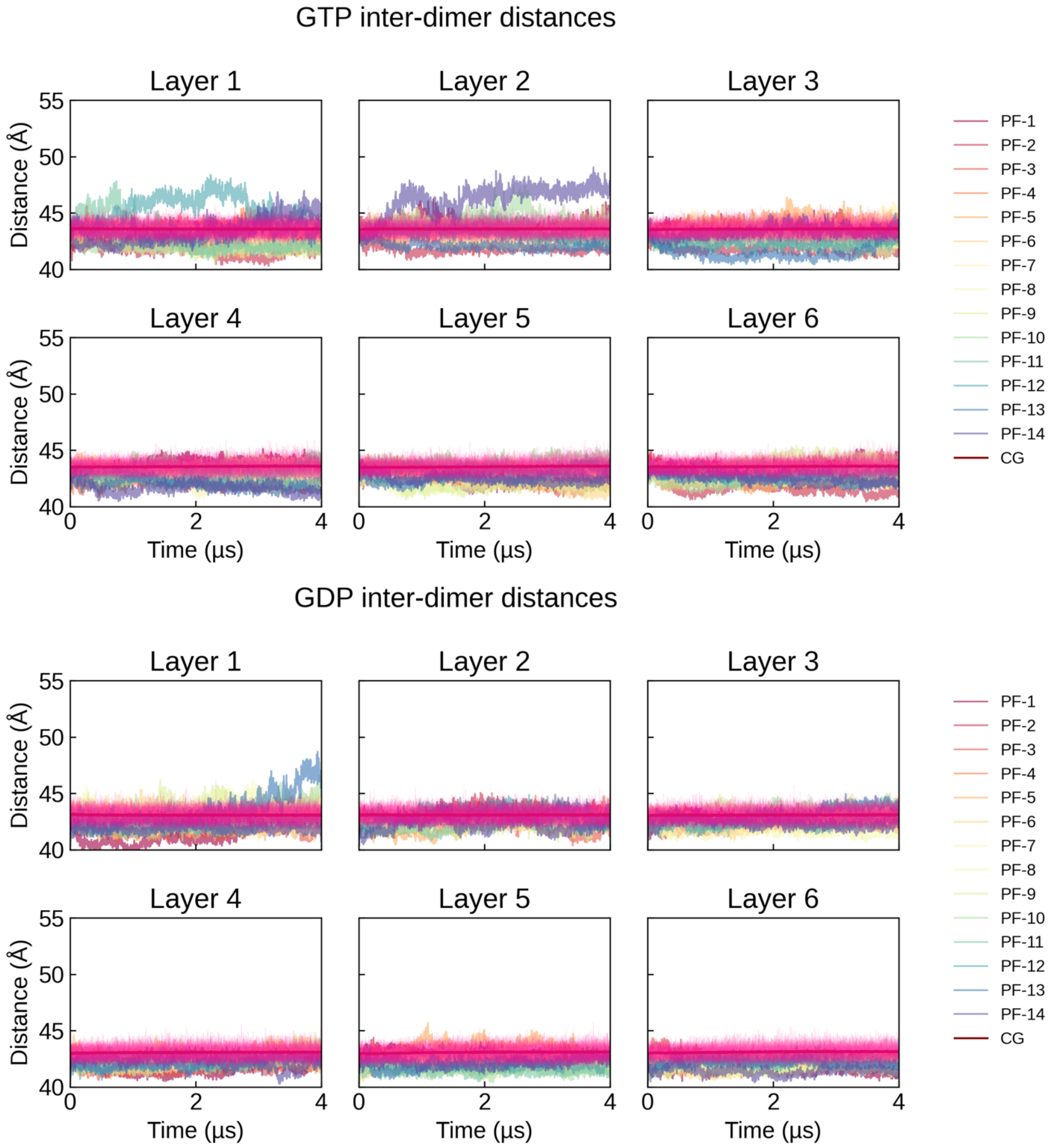

**Figure S1.11**: Snapshots of 40-layer simulations of GDP-MT with only 1 bottom layer restrained. This did not provide enough restraining force for the upper layers and have caused abnormal curvature of PFs near the boundary.

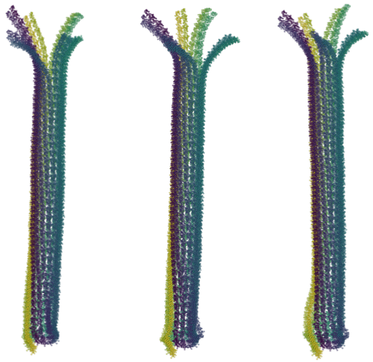

**Figure S1.12.** CG MD simulation snapshots on 40-layer 14-PF MT models. Snapshots taken at 20 µs. Simulations 1-9 are conducted with 2 layers of positional restraints at the bottom layers, and simulations 10-13 are conducted with 3 layers of positional restraints, also shown in Main Text Figure 6.

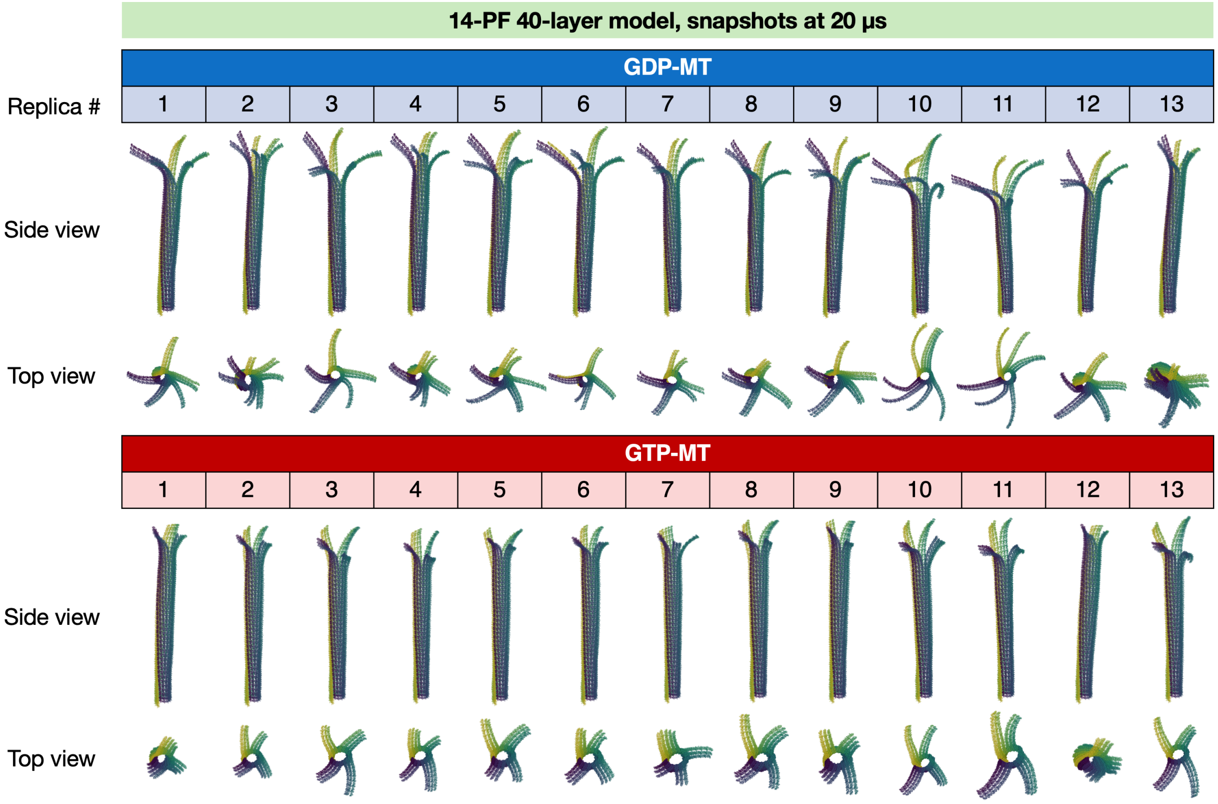

**Figure S1.13.** CG MD simulation snapshots on 40-layer 13-PF MT models. Snapshots taken at 20 μs.

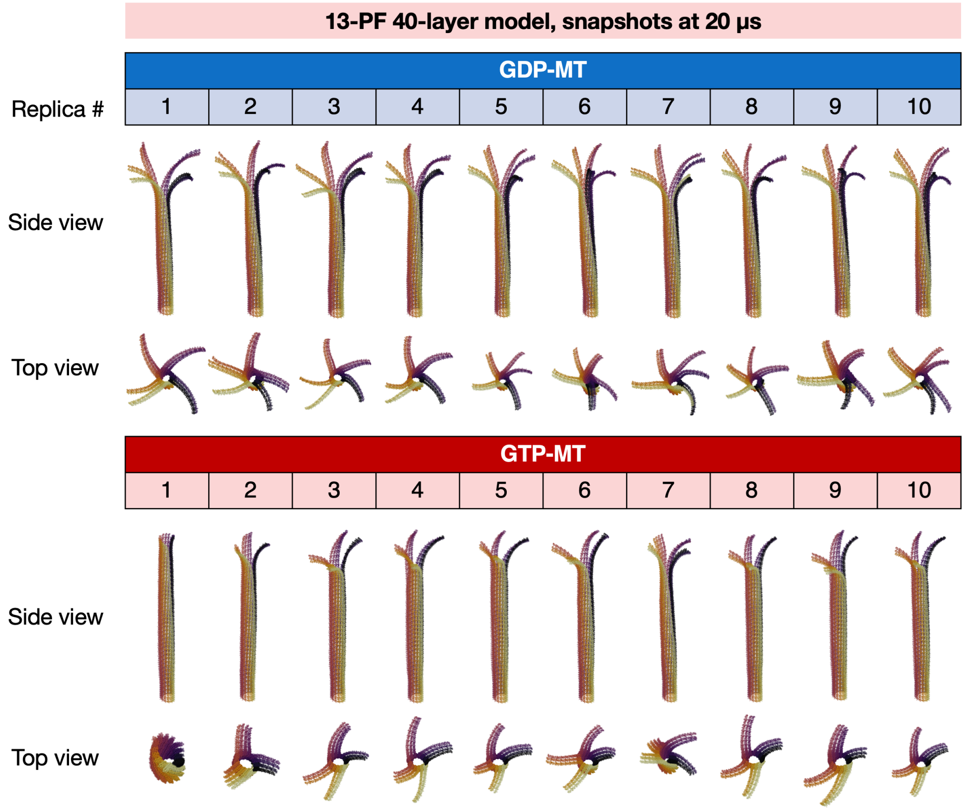


**Figure S1.14.** Statistics of 13-PF simulations as in Main Text Figure 7.

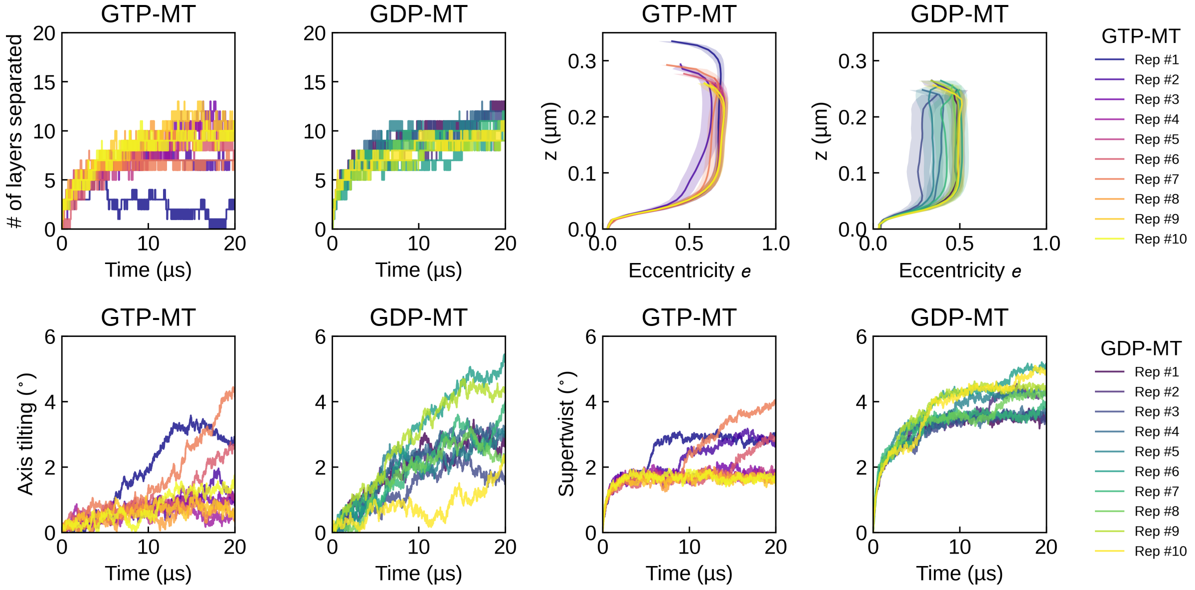

**Figure S1.15.** Outward peeling per protofilament for each 14-PF simulation. PF-$i$ means lateral dissociation of PF-$i$ and PF-$(i+1)$.

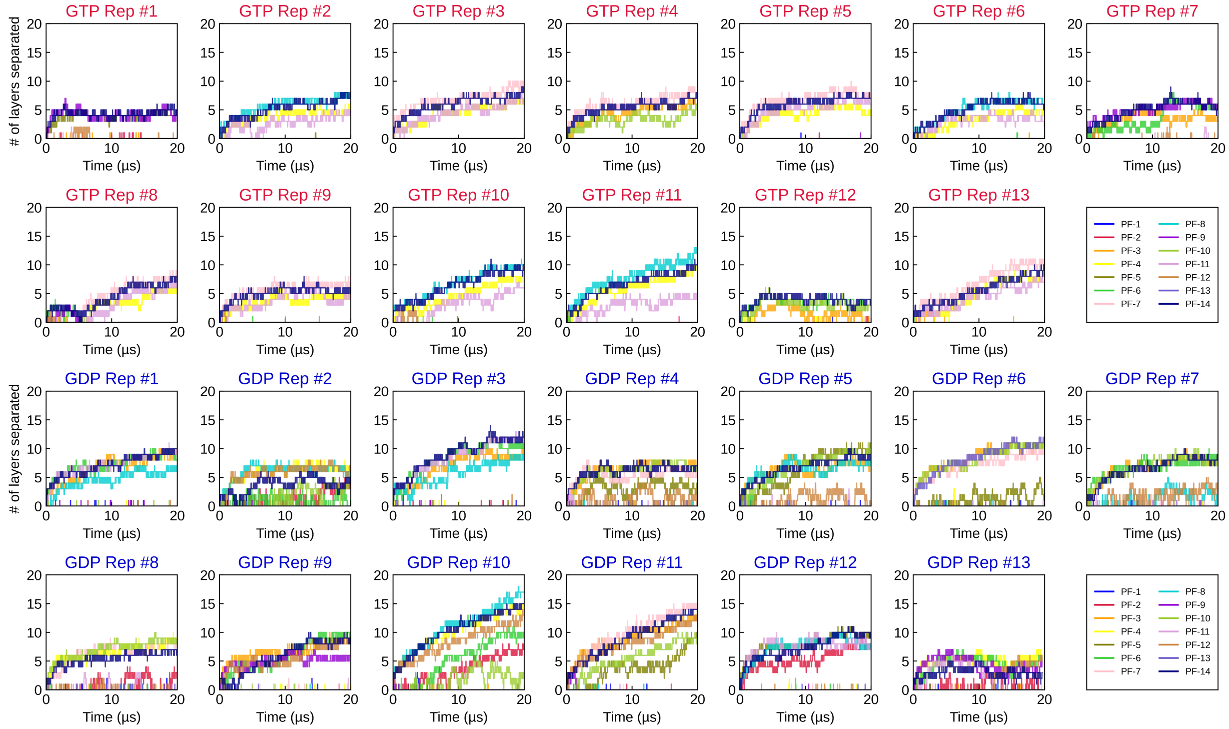


**Figure S1.16.** Outward peeling per protofilament for each 13-PF simulation. PF-$i$ means lateral dissociation of PF-$i$ and PF-$(i+1)$.

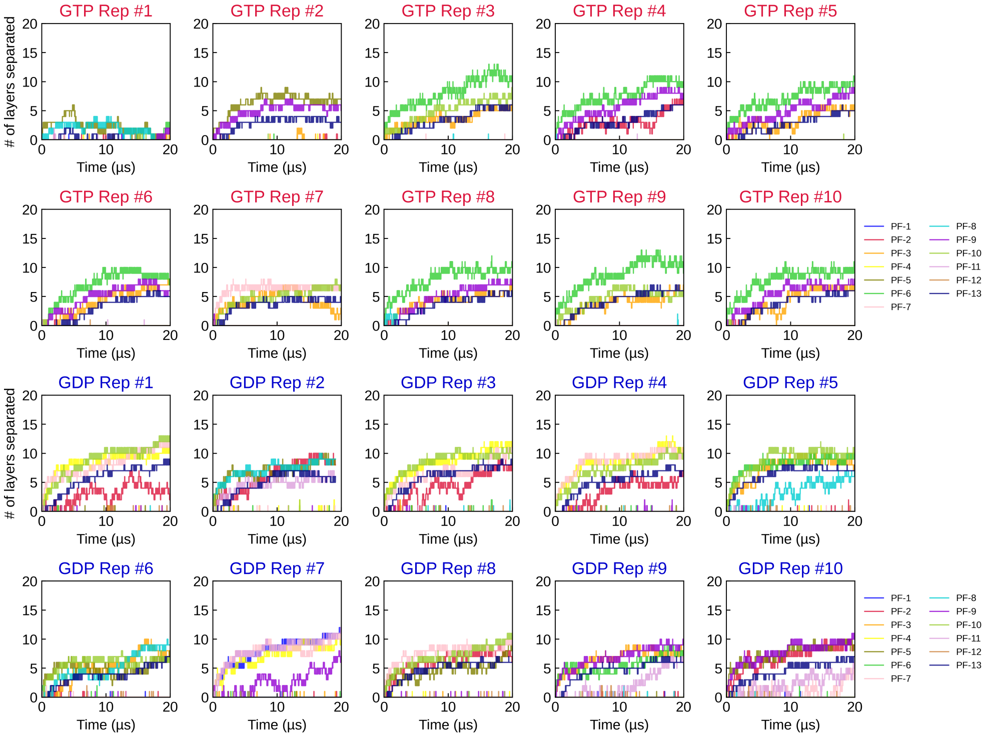

**Fig. S1.17.** The effect of added interlayer spacing in building long MT models. Here the example was done for 13-PF model. These are x-axis and y-axis projections of the MT lattice. We can see that especially for GDP-MT, a lattice spacing of 1.25 angstroms must be added to avoid lattice rupture.

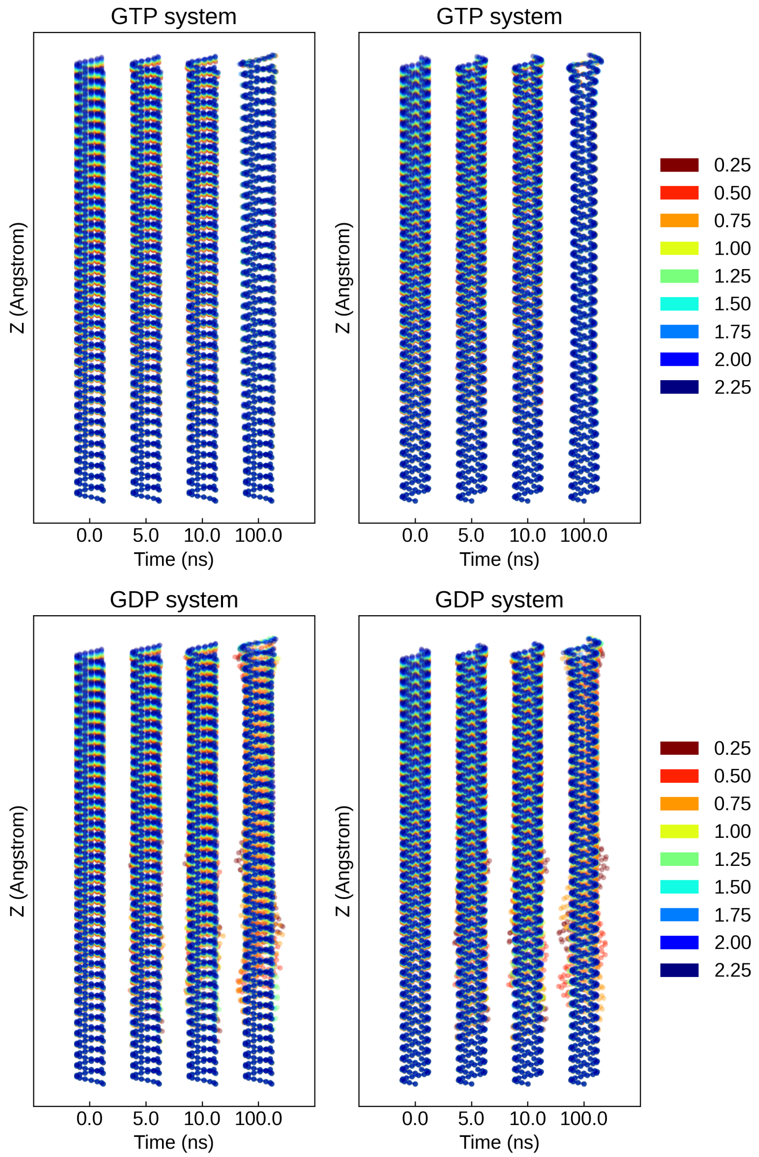


### Section 2: All-atom analysis

**Figure S2.1:** Average number of H-bonds in all lateral pairs, in different segments of simulation.

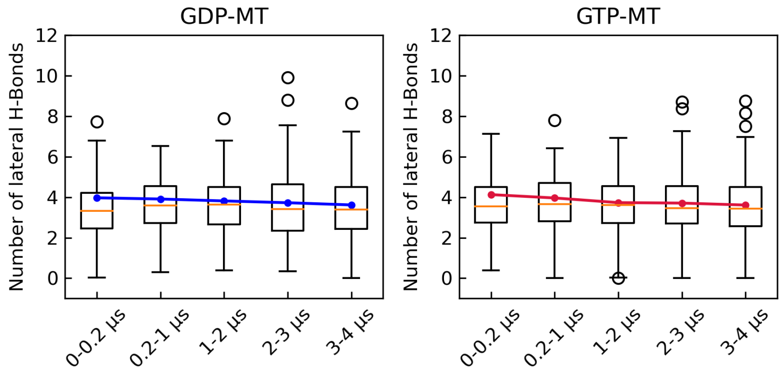


**Figure S2.2:** Average number of H-bonds in lateral pairs in the MT lattice, from 0 to 0.2 μs.

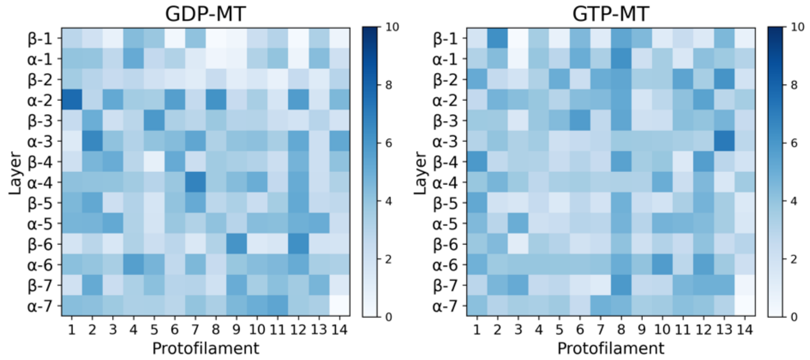


**Figure S2.3:** Average number of H-bonds in lateral pairs in the MT lattice, from 3 to 4 μs.

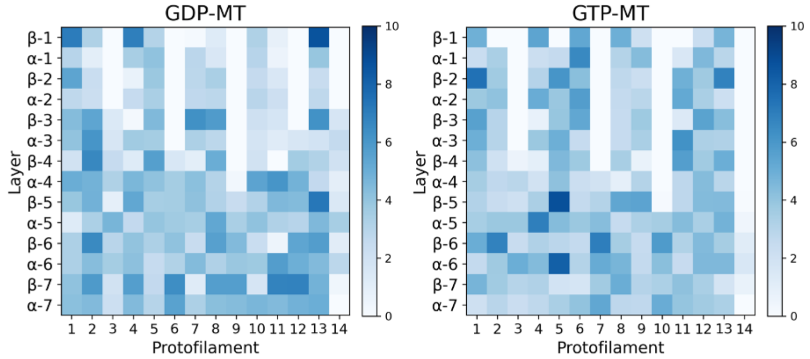

**Figure S2.4:** Frequency and correlation of the most frequent 12 H-bonds in the beta-beta lateral interface, averaged among different monomer pairs, in 0-1 μs and 3-4 μs segments of the trajectory.

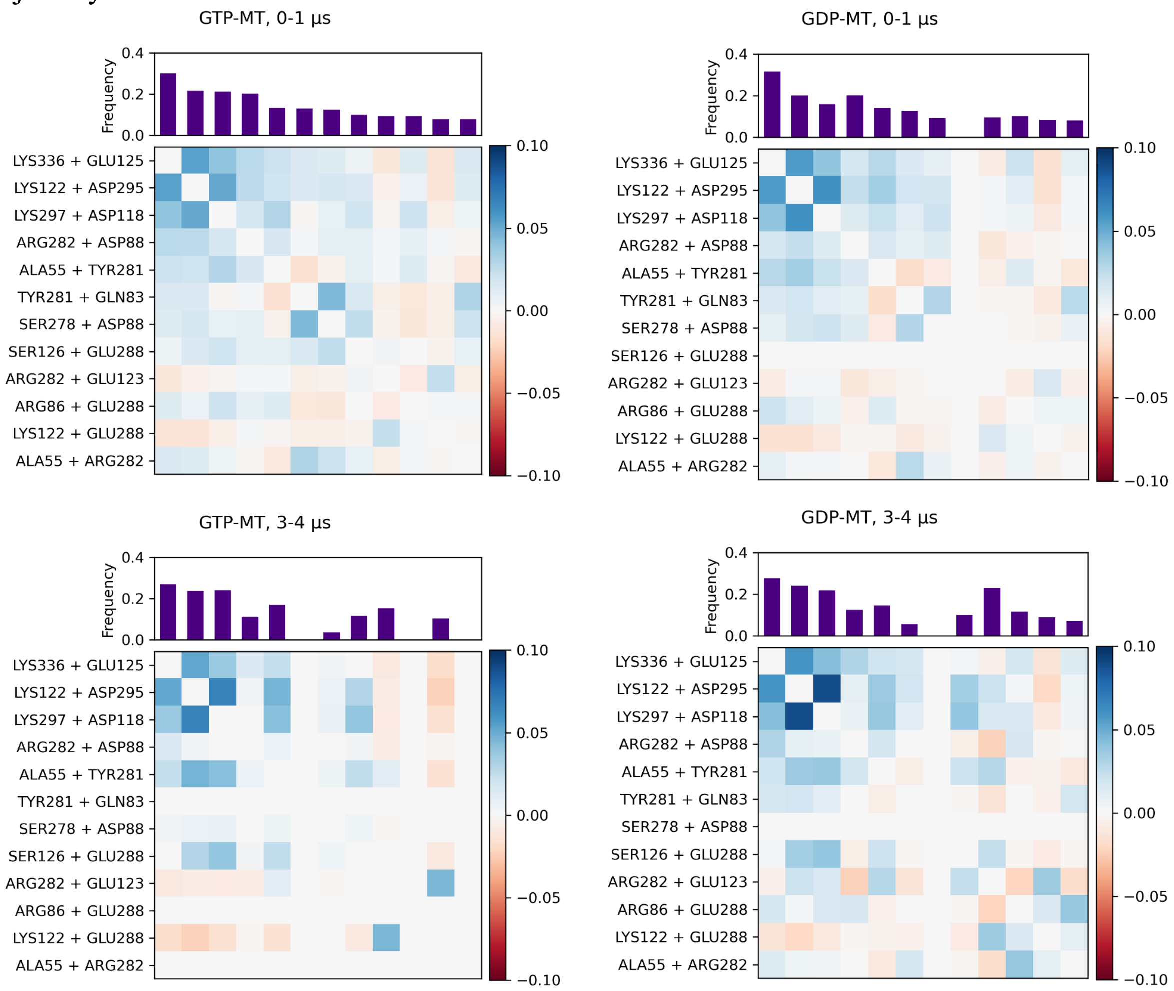

**Figure S2.5:** Frequency and correlation of the most frequent 12 H-bonds in the alpha-alpha lateral interface, averaged among different monomer pairs, in 0-1 μs and 3-4 μs segments of the trajectory.

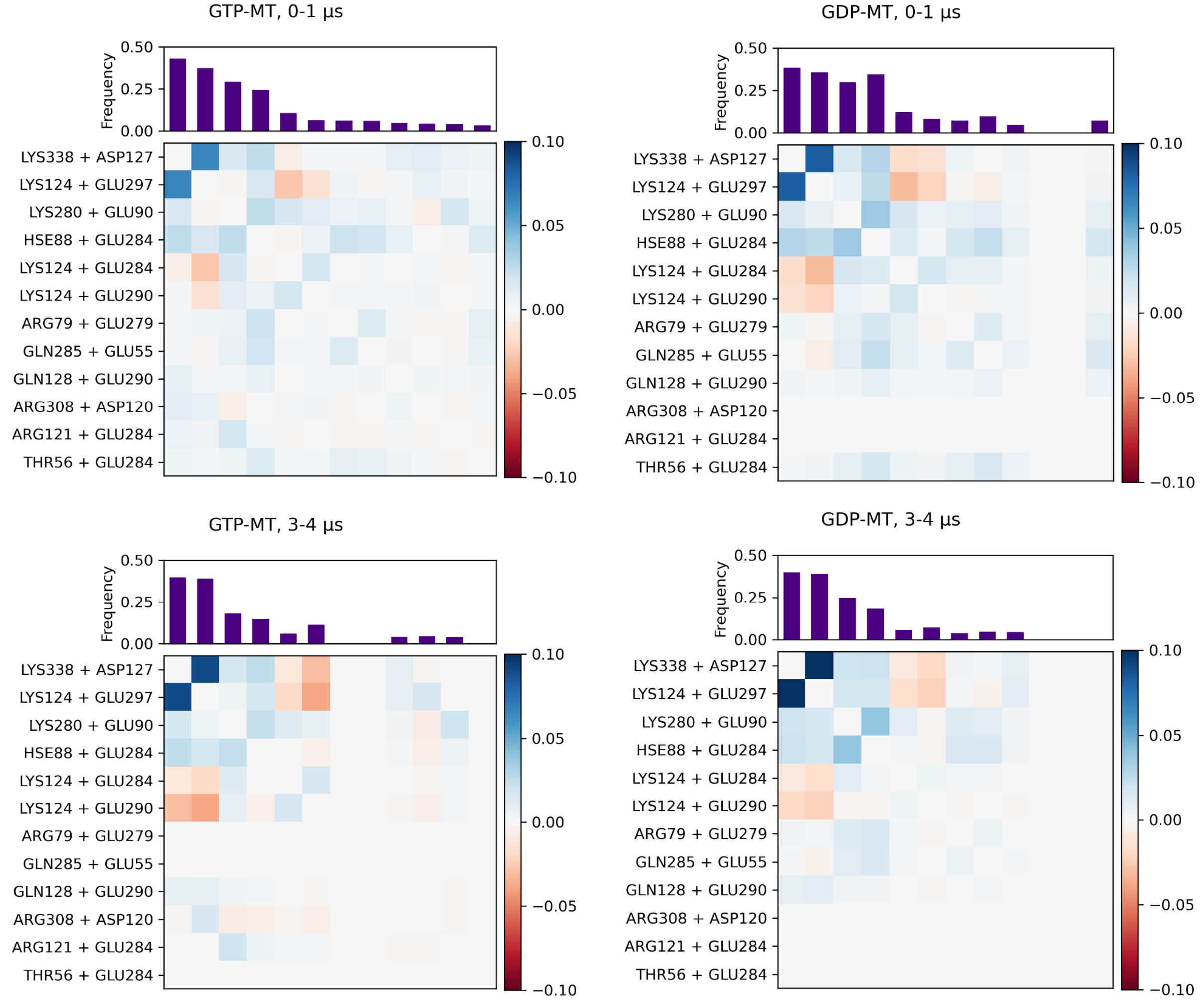

**Figure S2.6:** Frequency and correlation of the most frequent 12 H-bonds in the intra-dimer longitudinal interface, averaged among different heterodimers, in 0-1 μs and 3-4 μs segments of the trajectory.

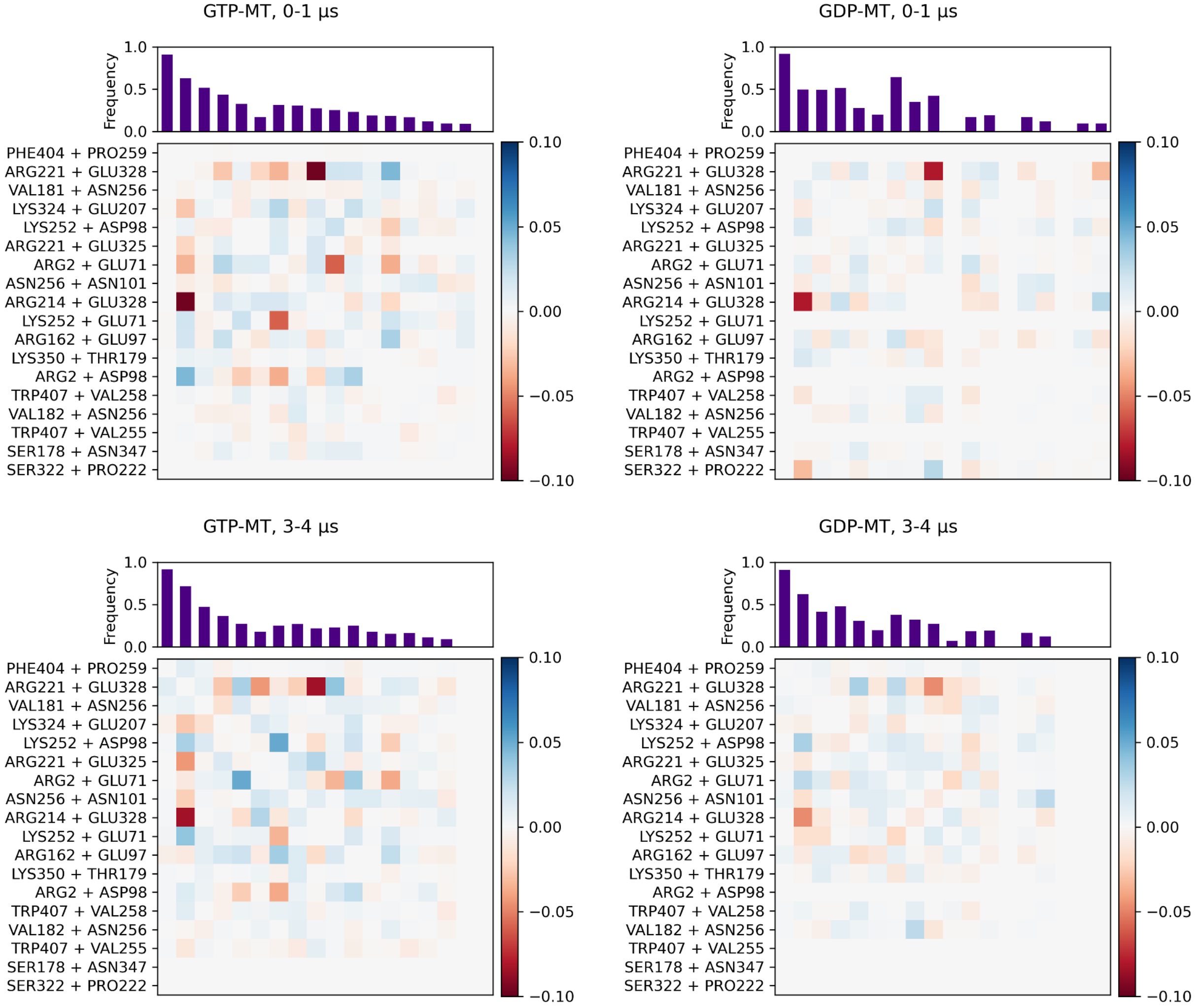

**Figure S2.7:** Frequency and correlation of the most frequent 12 H-bonds in the inter-dimer longitudinal interface, averaged among different longitudinal heterodimer pairs, in 0-1 μs and 3-4 μs segments of the trajectory.

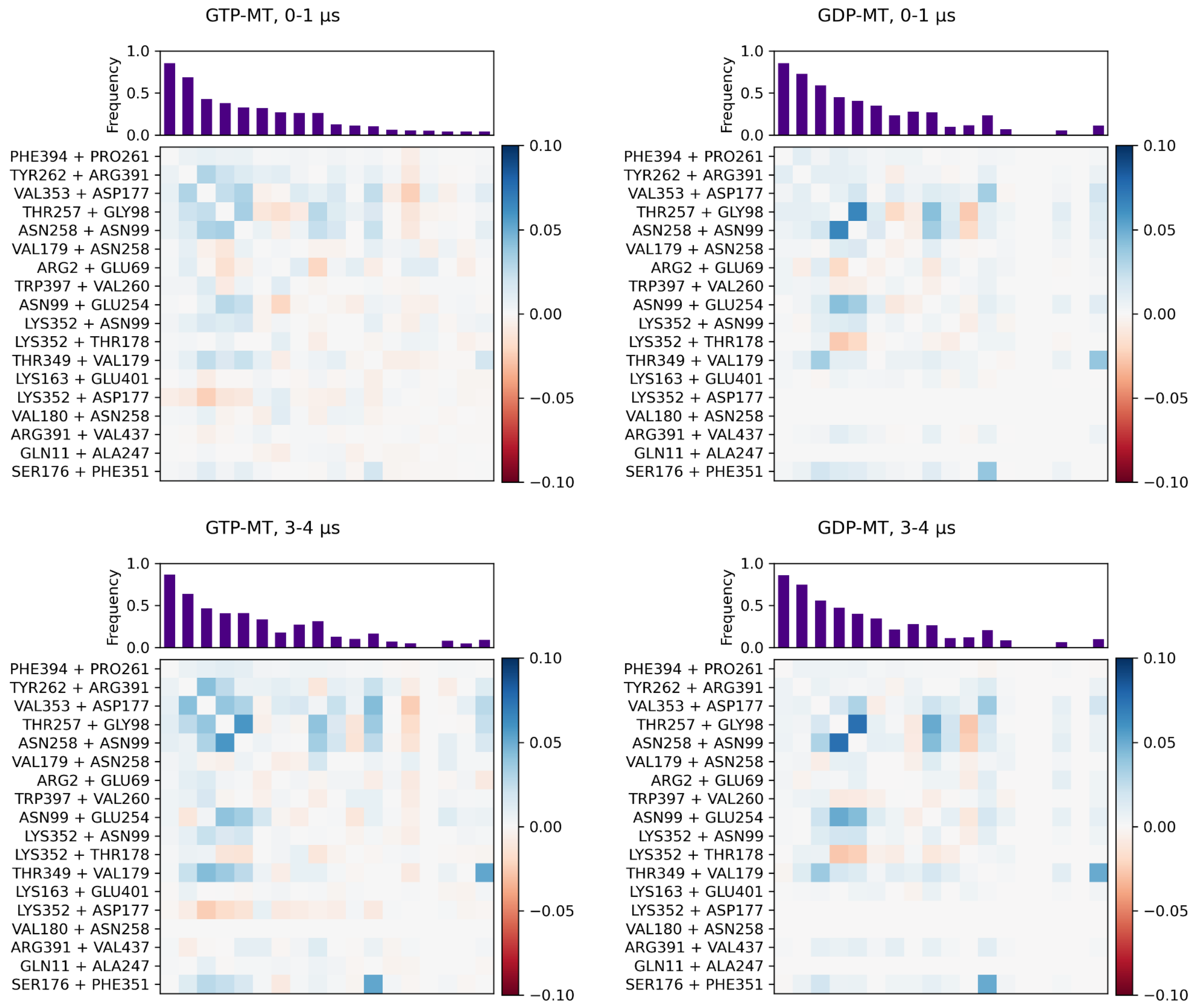

**Figure S2.8:** Comparison between the lateral interaction interfaces of alpha- and beta-tubulins. (a) Difference in H9-H3 interactions. (b) difference in M-loop interactions. (c) Arg-Arg stacking in beta-tubulin competes with H-bond formation. (d) Helical frequency of the M-loop, i.e., the fraction of frames in which the M-loop adopt alpha-helical structure as determined by the DSSP algorithm.

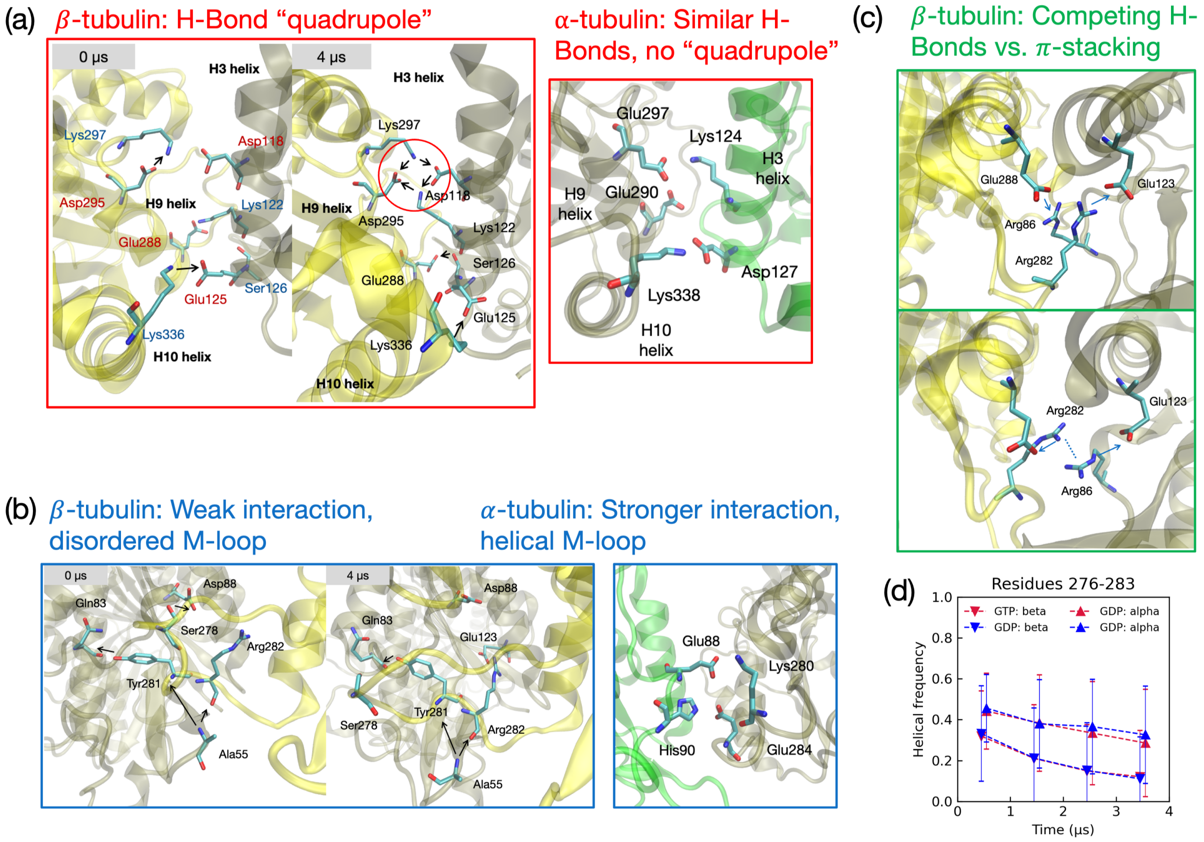

**Figure S2.9:** K-Means Clustering Scans on mixed distribution of GTP-MT and GDP-MT, and the projection on first two principal components (PCs) of data points with . The color coding corresponds to Figure 7c: states 0-3 corresponds to blue, red, purple and yellow, respectively.

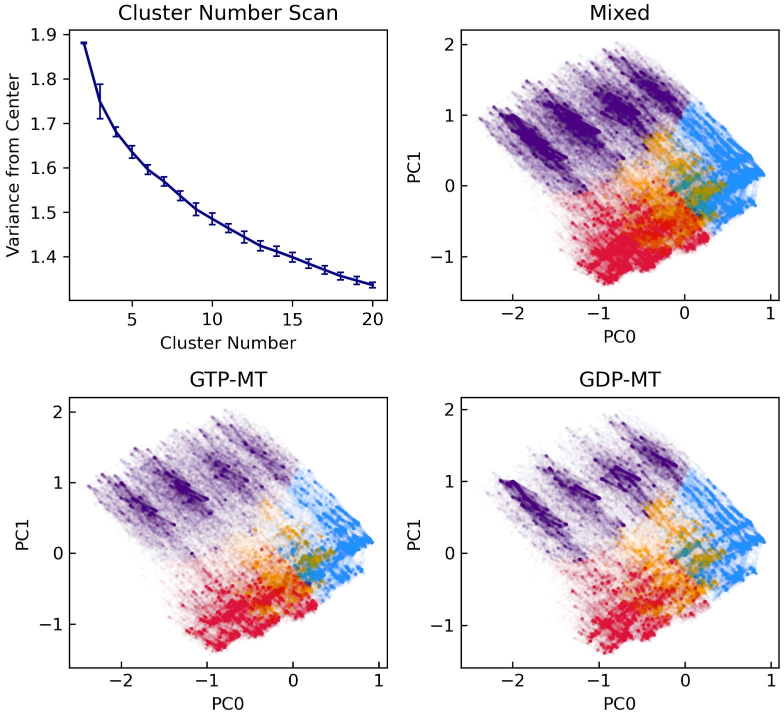

**Figure S2.10:** Probabilities of lateral interaction Mode 0 across the MT lattice, in 4 segments of the 4 μs all-atom trajectory.

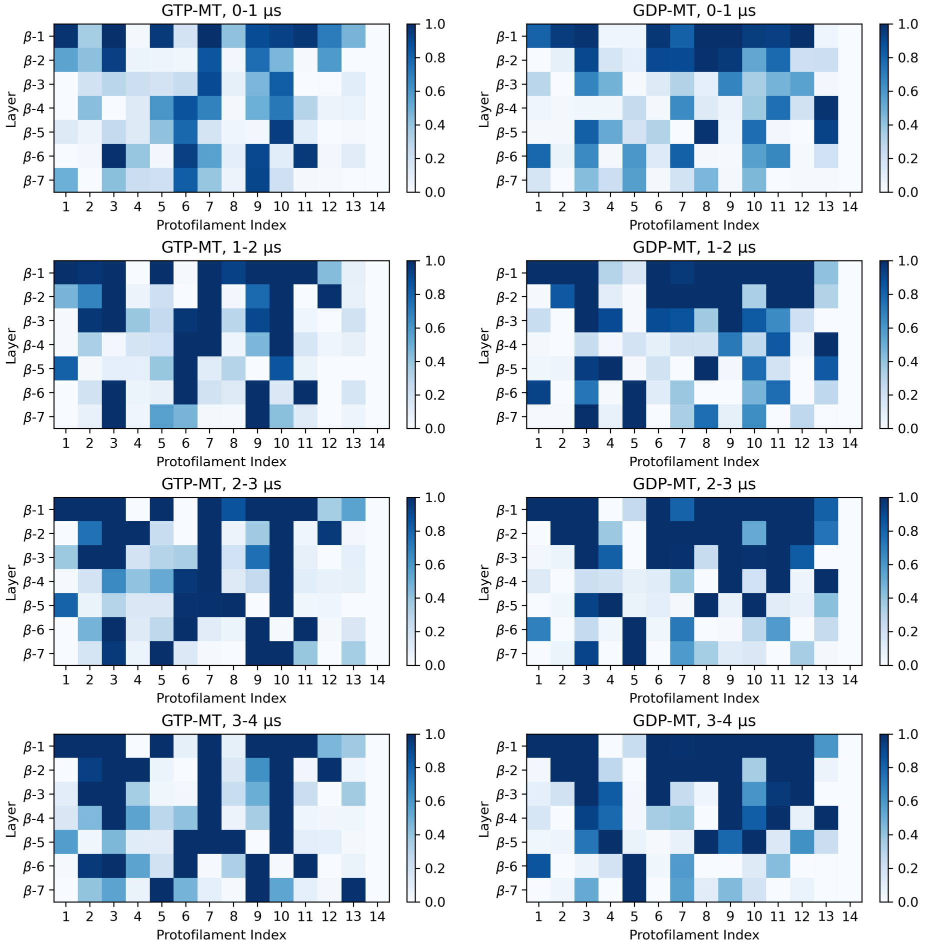

**Figure S2.11:** Probabilities of lateral interaction Mode 1 across the MT lattice, in 4 segments of the 4 μs all-atom trajectory. Mode 1 increases in frequency as the MT system relaxes.

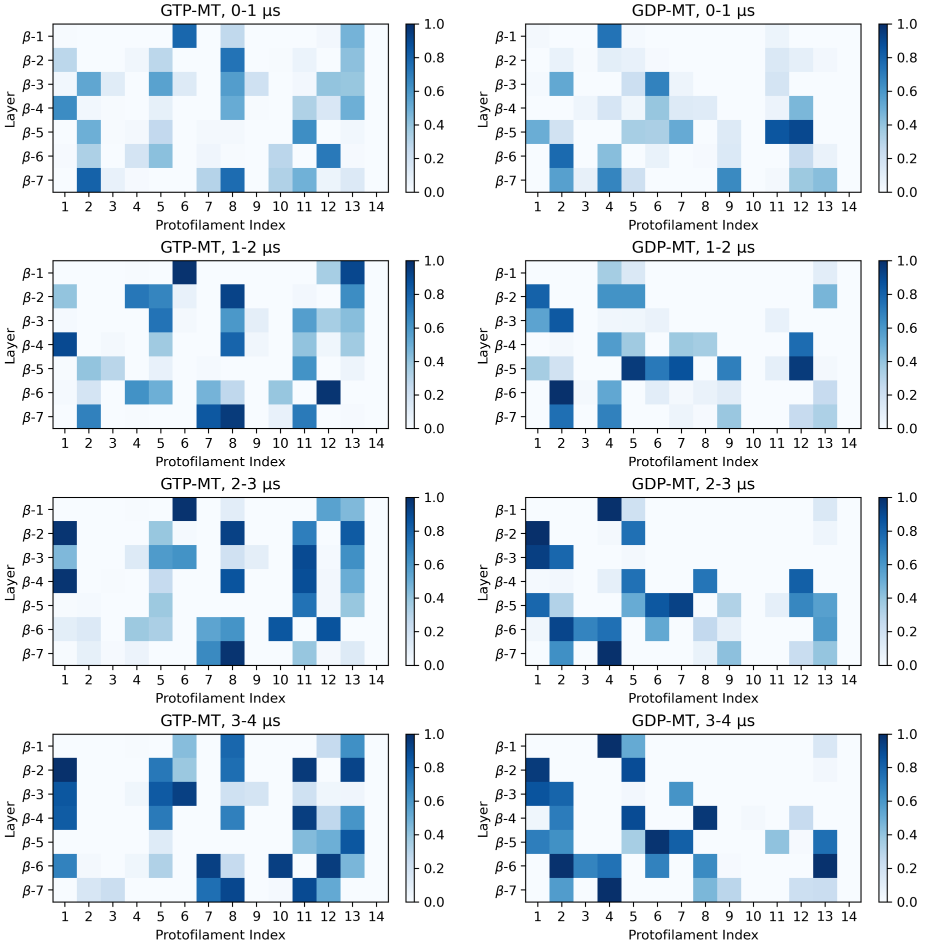

**Figure S2.12:** Probabilities of lateral interaction Mode 2 across the MT lattice, in 4 segments of the 4 μs all-atom trajectory. Mode 2 decreases in frequency as the MT system relaxes.

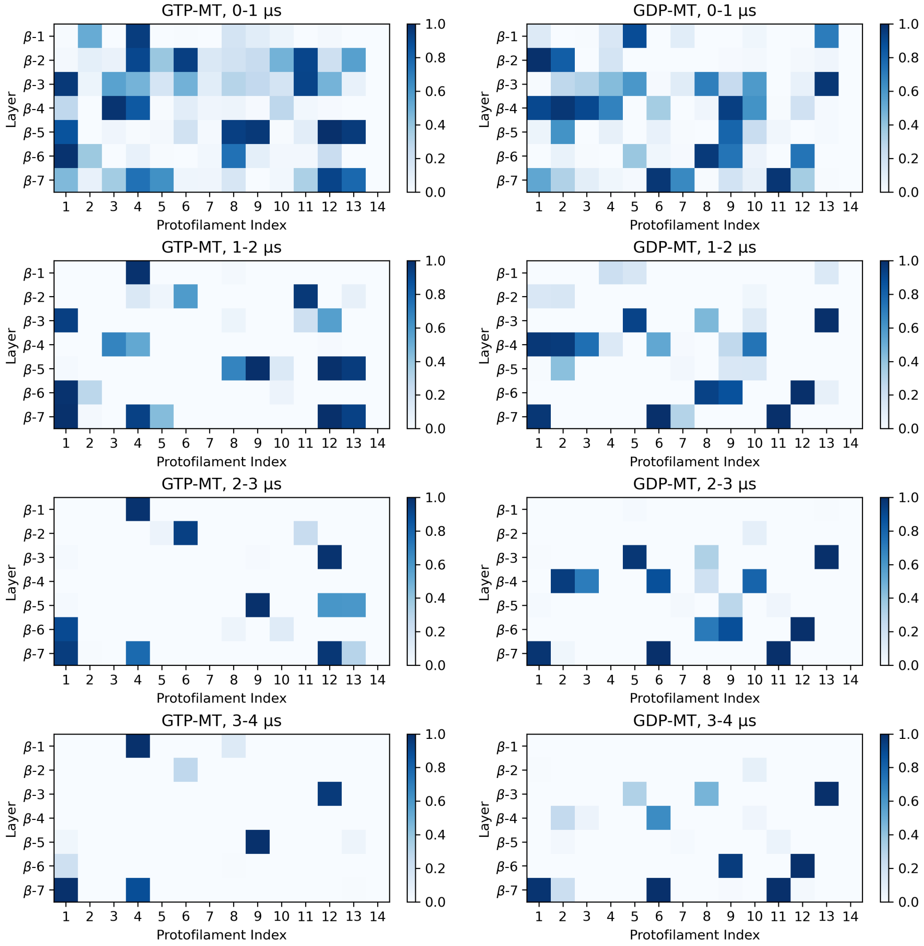

**Figure S2.13**: Probabilities of lateral interaction Mode 3 across the MT lattice, in 4 segments of the 4 μs all-atom trajectory.

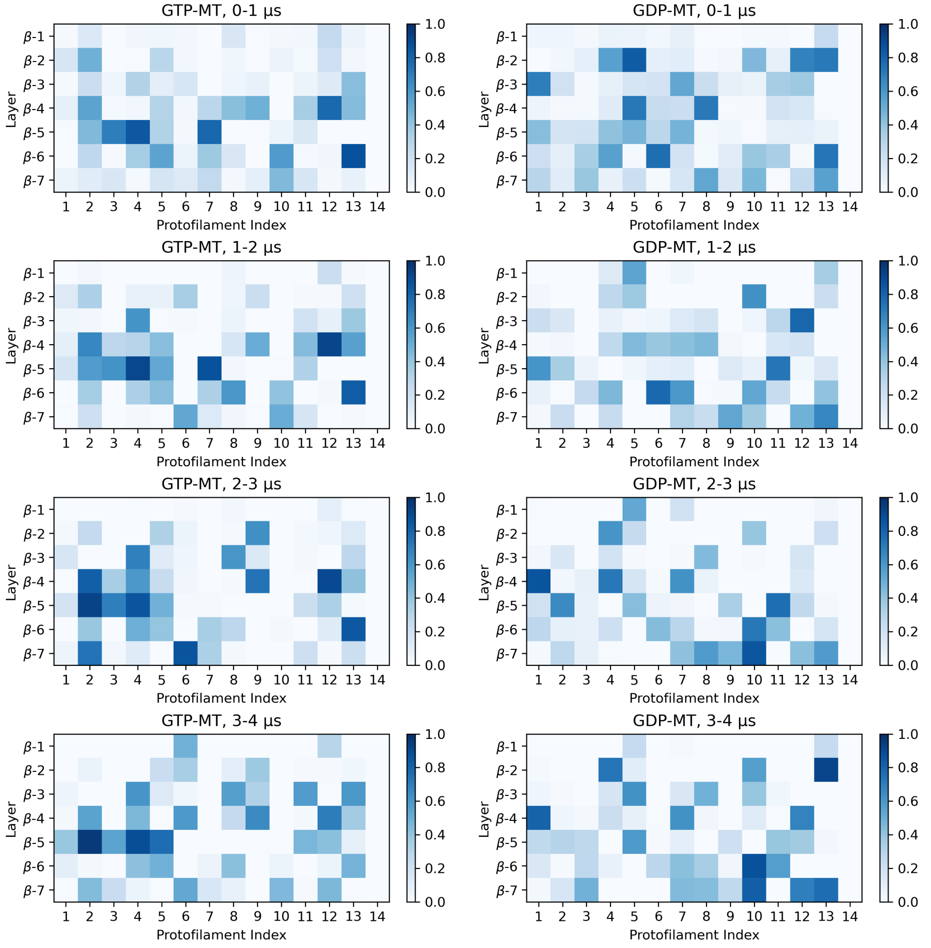

**Figure S2.14:** Time evolution of the mean H9-H3 dihedral angle in alpha-tubulin (See Figure 7d for beta-tubulin) in each heterodimer layer. Dissociated pairs excluded to remove noise.

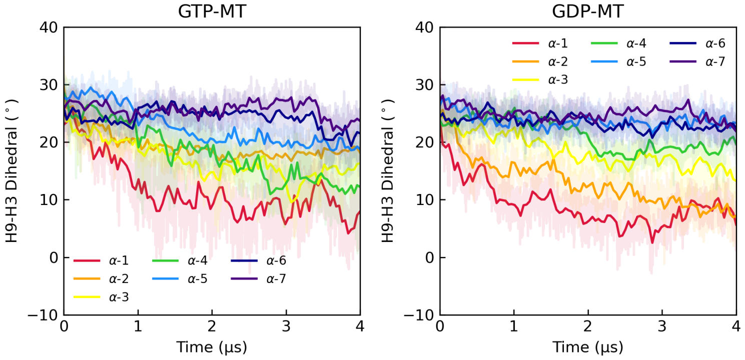


**Figure S2.15:** H9-H3 helix distance and crossing angle distribution of beta-tubulin in each heterodimer layer, with comparison between 0-0.1 µs and 0.5-0.6 µs. The distance distribution is bimodal and shifted in the left peak's favor in 0.5 µs. Dissociated pairs excluded to remove noise.

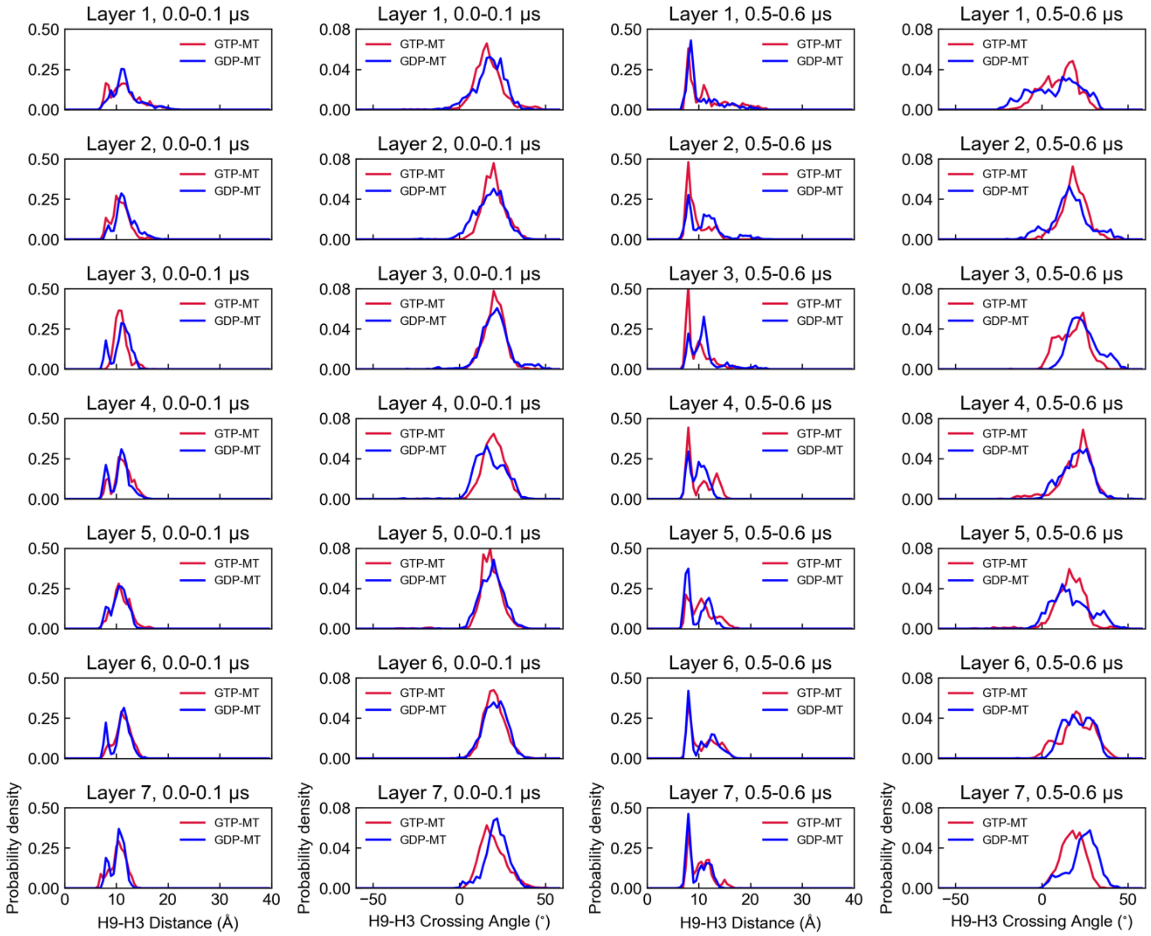

**Figure S2.16:** H9-H3 helix distance and crossing angle distribution of beta-tubulin in each heterodimer layer, with comparison between 0-1 μs and 3-4 μs.

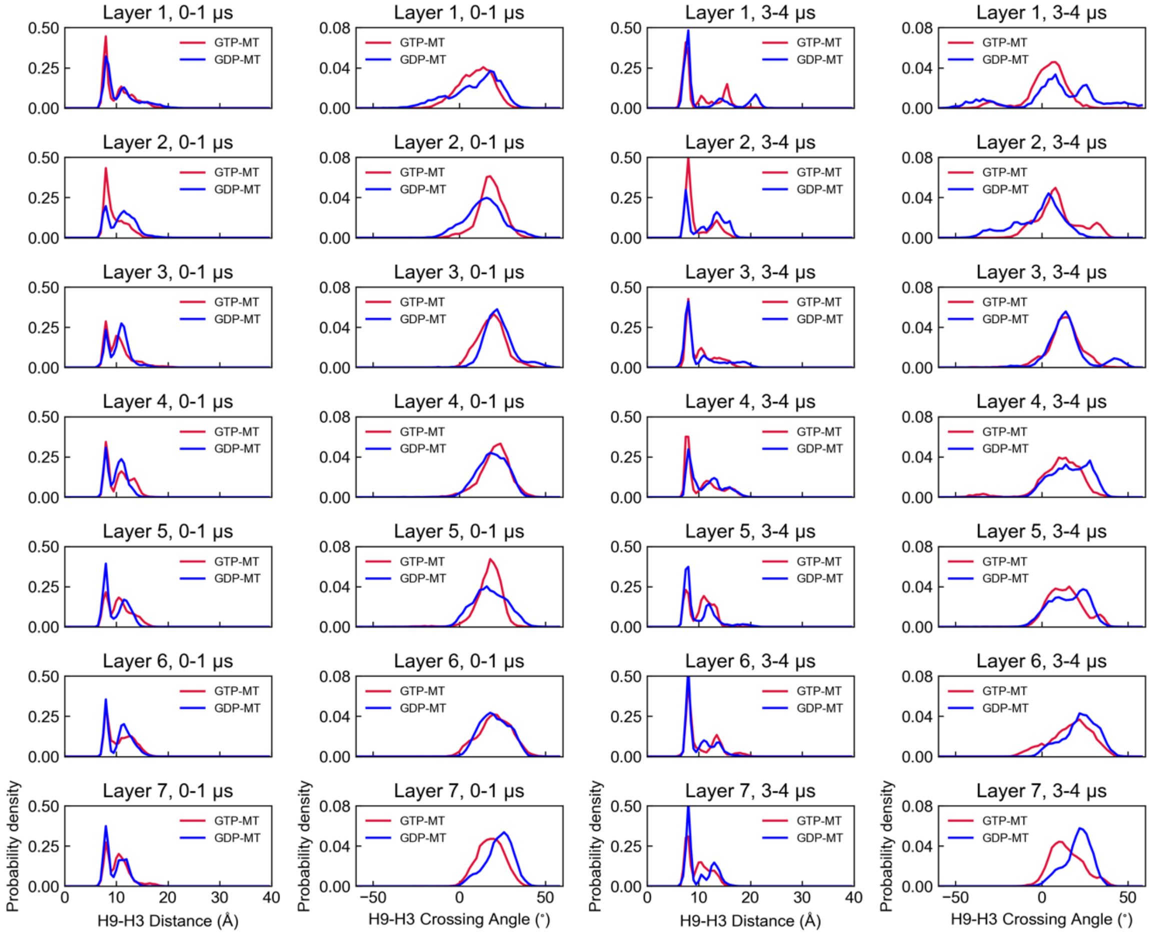

**Figure S2.17:** H9-H3 helix distance and crossing angle distribution of alpha-tubulin in each heterodimer layer, with comparison between 0-0.1 μs and 0.5-0.6 μs.

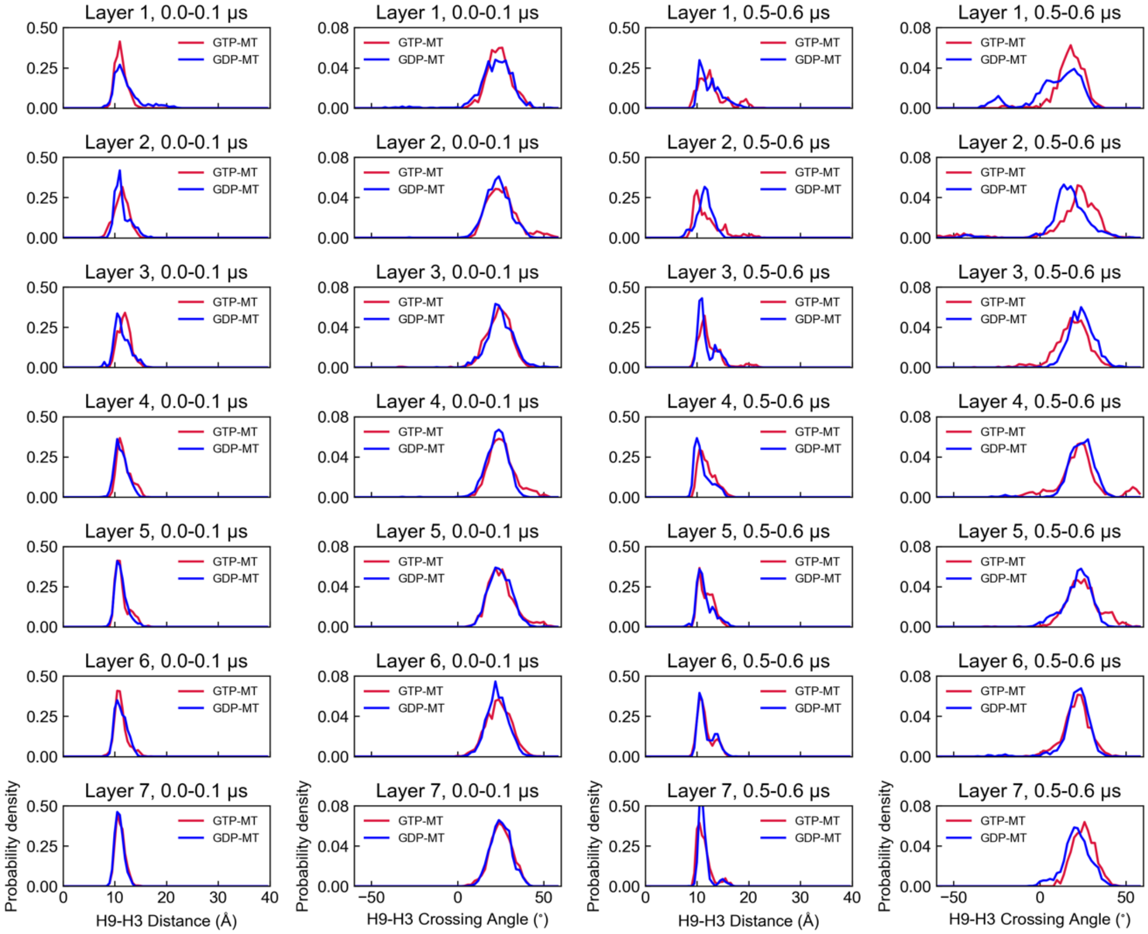

**Figure S2.18:** H9-H3 helix distance and crossing angle distribution of alpha-tubulin in each heterodimer layer, with comparison between 0-1 μs and 3-4 μs.

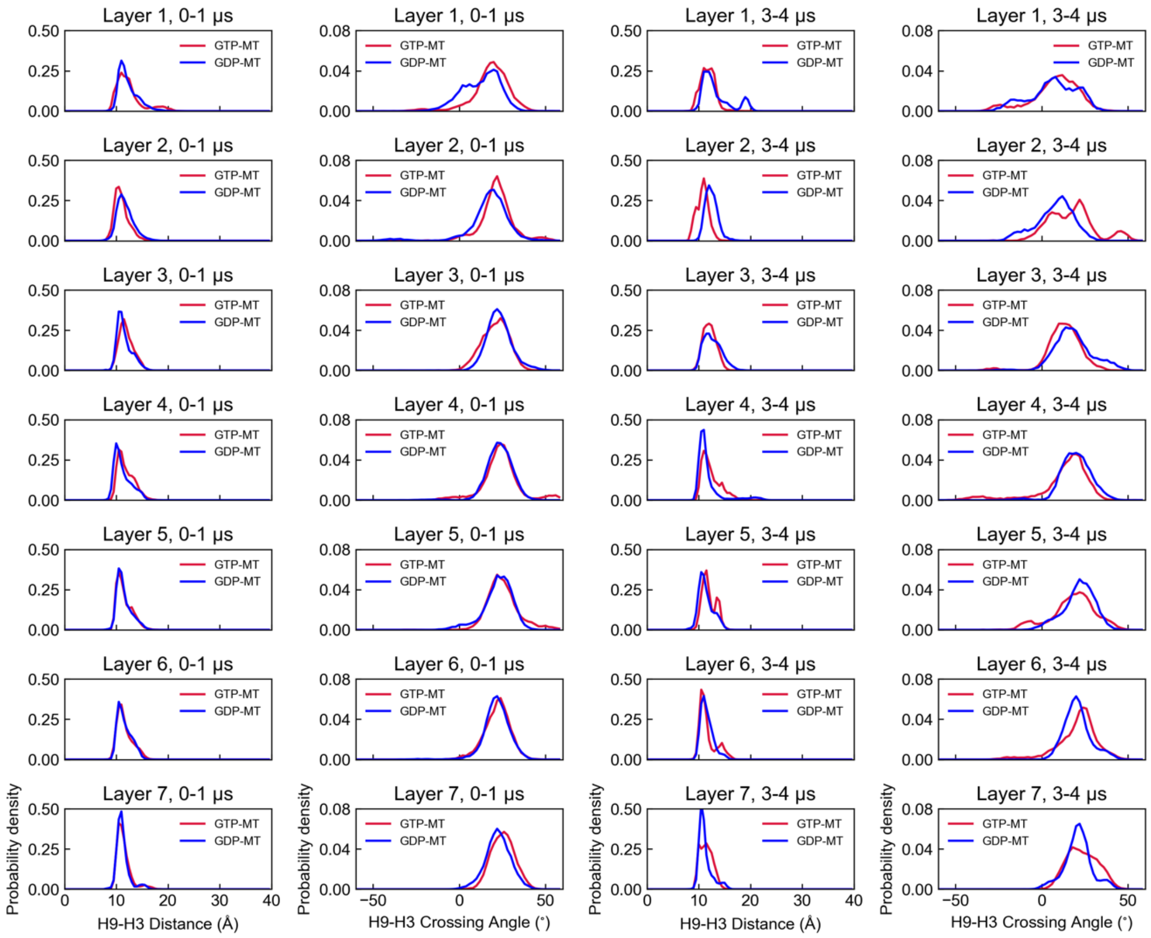

**Figure S2.19:** Time evolution of the H9-H3 distance between the eventually dissociated PF pairs at the top heterodimer layer (Layer 1).

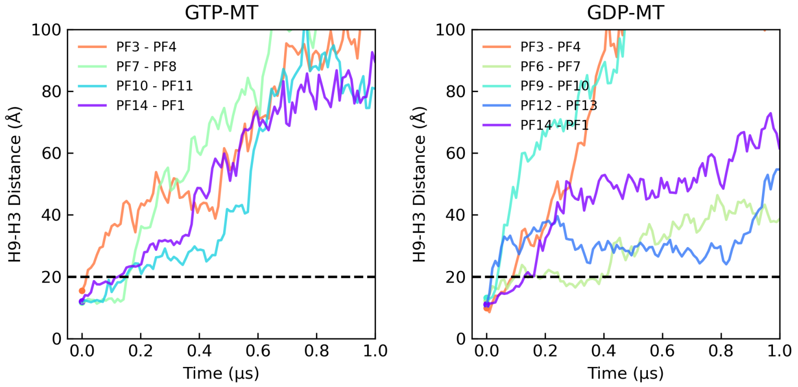


**Figure S2.20:** Time evolution of average inter-dimer COM distance in each heterodimer layer.

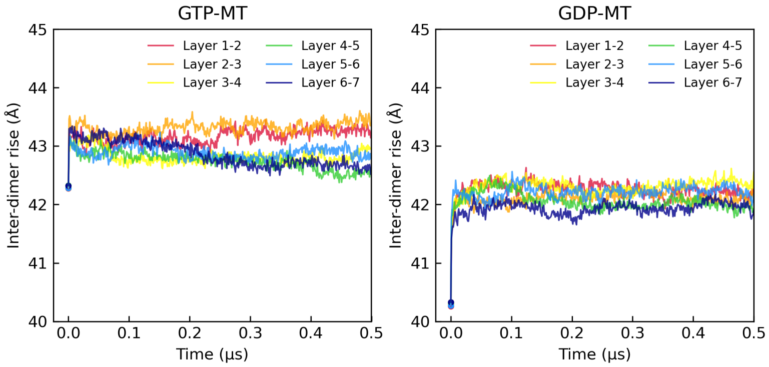


**Figure S2.21:** Illustration of H10-H6 H-bonds, including β:Lys324-α:Glu207 and β:Glu325/328-α:Arg221.

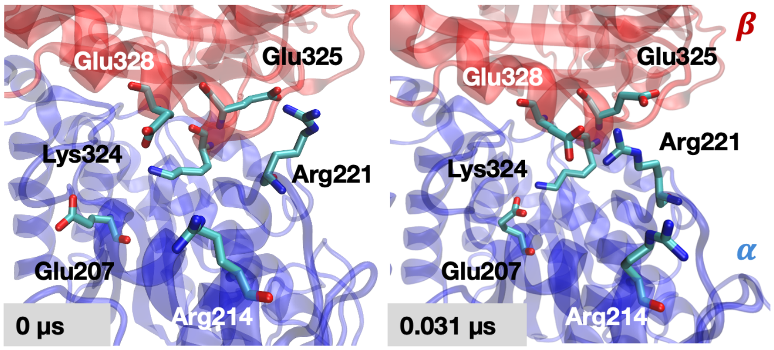

**Figure S2.22:** Time evolution of average Cα-Cα and N-O distance in the β:Lys324-α:Glu207 H-bond pair, synchronous to the intra-dimer distance jump.

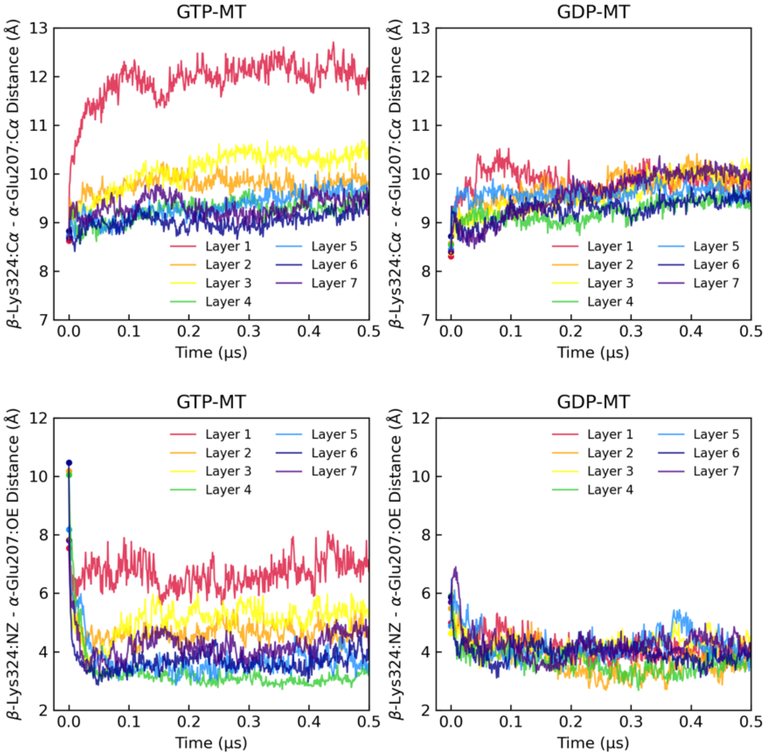


**Figure S2.23:** The direction of initial perturbation at Layer 1, characterized by the trajectories of $\Delta r_i = r_{1,i} - r_{2,i}$, $\Delta z_i = z_{1,i} - z_{2,i}$, $\Delta\phi_i = \phi_{1,i} - \phi_{2,i}$, averaged for the left, middle and right PFs in the PF clusters. Notice the swift increase in both $z$ and $r$ components.

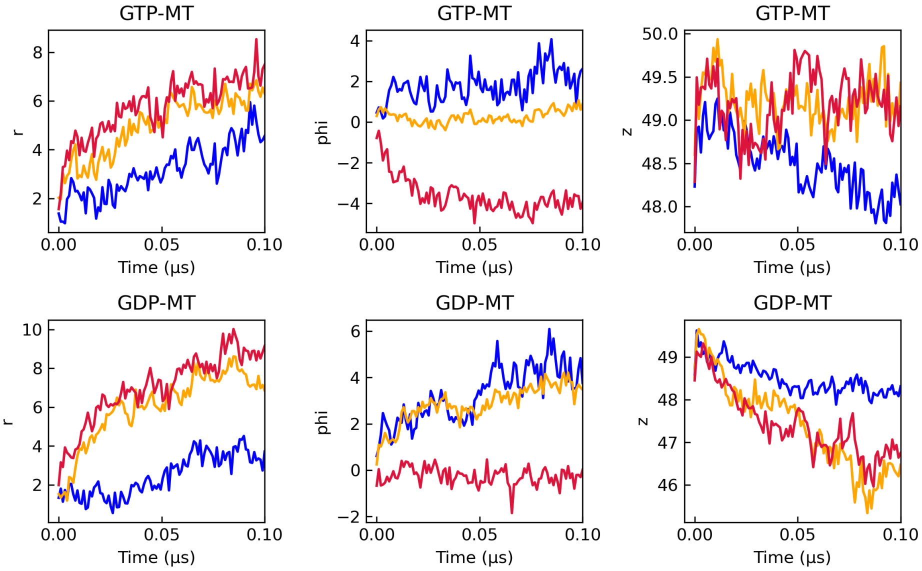
